# Rational antigen engineering and mucosal delivery design for next-generation RSV vaccines

**DOI:** 10.64898/2026.07.28.739720

**Authors:** Wangjun Fu, Desheng Liu, Xiangfei Shan, Chujun Ding, Hui Zhai, Lu Jiang, Yuqing Zhou, Wenqiang Mao, Jie Deng, Mingkai Li, Yaling Hu, Zhe Lv, Yufei Xia, Xiangxi Wang

**Author notes:** These authors contributed equally to this work (Wangjun Fu, Desheng Liu, Xiangfei Shan, Chujun Ding). Correspondence (Z. L); (YF. X); (XX.W).

## Abstract

Respiratory syncytial virus (RSV) prefusion F (preF) vaccines have transformed adult prophylaxis, yet unmet needs in antigen stability, pediatric safety, and mucosal protection persist. Here, we develop an integrated structure-guided RSV vaccine design platform that couples allosteric stabilization, epitope-focused immunogen engineering, and route-specific mRNA delivery for systemic and mucosal immune activations. By mapping prefusion F “breathing” motions and applying a ThermoNet-and Rosetta-guided screening funnel, we identified R296, a stabilized prefusion F immunogen that reinforces the α1–α5 hinge and interprotomer interfaces while preserving key neutralizing epitopes. Cryo-EM confirmed that R296 retains a native-like prefusion architecture. And mRNA-LNP vaccination elicited potent, durable, and broadly protective neutralizing responses in mice, rats, and cotton rats, with clearance of detectable infectious virus and no evidence of Th2-skewed enhanced respiratory disease. To address pediatric safety, we designed a stalkless nanoparticle immunogen, Head38-50AB-3, which enriches high-potency apical epitopes while excluding stalk regions associated with low-potency or non-protective responses, conferring protection without VAERD-like pathology. Finally, we engineered an intranasal-delivered LNP that enables intranasal R296 mRNA delivery, inducing systemic neutralization together with robust nasal and bronchoalveolar secretory IgA (sIgA). R296 has now advanced to Phase 1 clinical trials. These results establish a modular framework for next-generation RSV vaccines.

## Introduction

Respiratory syncytial virus (RSV) is a ubiquitous and highly contagious pathogen that imposes a substantial global burden of acute lower respiratory tract infections, particularly in infants, older adults, and immunocompromised populations^1-3^. The past three years have marked a turning point for RSV intervention, with regulatory approvals granted for three prefusion F (preF) glycoprotein-based vaccines for older adults: GSK’s Arexvy, Pfizer’s Abrysvo, and Moderna’s mRESVIA^4-6^. These vaccines leverage the structural insight that the metastable preF conformation presents key neutralizing epitopes (such as site Ø) that are conformationally rearranged in the postfusion state^7-9^. While these advances support the preF as a primary vaccine target, the field continues to grapple with limitations in antigen design, durability, and safety profiles across different age groups^10-14^. Current preF-stabilization strategies, beginning with the DS-Cav1 scaffold, have largely relied on engineered disulfide bonds and cavity-filling substitutions, and later designs further incorporated proline substitutions to rigidify metastable regions of RSV F^15-18^. However, the structural plasticity of the F protein suggests that alternative allosteric vulnerabilities exist. The identification of additional, energetically rational stabilization strategies that reinforce interprotomer interfaces and hinge regions remains a critical frontier for improving thermostability and manufacturability^16,19-21^.

Beyond the need for improved structural stabilization, significant gaps remain in protective efficacy and safety. Current licensed vaccines for older adults, while effective, exhibit waning immunity within months, necessitating ongoing research into more durable immunogens or regimens^10,11^. Furthermore, systemic administration fails to elicit robust mucosal immunity at the primary site of infection^22^. This lack of durable secretory IgA (sIgA) and tissue-resident memory T cells leaves individuals susceptible to repeat infections and offers incomplete protection against viral transmission. The most pressing challenge, however, lies in pediatric applications^23^. Recent clinical holds on leading mRNA RSV vaccine candidates for infants, following observations of a numerical imbalance in severe lower respiratory tract illness in 5- to 8-month-olds, have underscored the fragility of the safety window^12^. These setbacks are hypothesized to stem from non-neutralizing or poorly neutralizing epitopes (e.g., stalk regions) that may drive Th2-biased immune responses and vaccine-associated enhanced respiratory disease (VAERD) ^24-28^, highlighting an urgent need for epitope-focused designs that exclude these domains.

The complexity of RSV vaccine development is further compounded by the antigenic landscape of the F protein^7,29^. Structural analyses reveal a dichotomy between potent, site-specific neutralizing antibodies targeted by the adult immune system and the lower-potency, germline-encoded responses prevalent in toddlers^30^. This suggests that stabilizing the protein alone is insufficient; the epitopes presented must be carefully curated to avoid priming non-protective immunity. Moreover, the route of administration plays a pivotal role in safety and efficacy. Intramuscular vaccines induce systemic IgG but fail to elicit the mucosal barrier immunity needed for effective protection. Conversely, intranasal delivery has historically faced challenges with rapid mucociliary clearance and inefficient transfection of respiratory epithelium, often failing to match the immunogenicity of parenteral routes without risking toxicity or instability^22,31^.

To address these multifaceted challenges, we developed a modular platform of structure-guided innovations targeting distinct unmet needs in RSV prophylaxis. First, we moved beyond empirical stabilization by computationally mapping the allosteric "breathing" motions of the F protein. Using a four-strategy screening funnel combining ThermoNet^32^ predictions, Rosetta^33^ scans, and conformational monitoring, we engineered a lead candidate, R296, that reinforces the α1–α5 hinge and interprotomer contacts without compromising antigenicity. Second, recognizing the pediatric safety imperative, we developed an epitope-guided, stalkless nanoparticle immunogen (Head38-50AB-3) that selectively displays high-potency apical epitopes while eliminating stalk regions putatively associated with historical VAERD risk. Third, to bridge the mucosal immunity gap, we engineered a nasal-delivery lipid nanoparticle that prolongs mucosal residence and enhances intranasal mRNA expression, thereby eliciting potent systemic cellular and humoral immunity together with sIgA responses in both nasal and lung mucosa. Together, these results provide a multi-pronged solution for next-generation RSV vaccines, offering enhanced stability, pediatric safety, and mucosal protection through rational design.

## Results

### Computational mapping of allosteric vulnerability and four-strategy screening funnel for prefusion F stabilization

During intracellular maturation, the RSV F precursor F0 is cleaved at two furin sites, releasing the intervening p27 peptide and generating a disulfide-linked F2–F1 heterodimer with the hydrophobic fusion peptide exposed at the N terminus of F1^34^. Triggering drives large-scale refolding: Refolding Region 1 (RR1, residues 137–216) collapses into the postfusion α5 helix, while Refolding Region 2 (RR2, near the C terminus of F1) swings to close the six-helix bundle (6HB) ^9,35^. This global refolding is coupled to localized, lower-energy motions—most notably a breathing between helix α1 (and the adjacent β2–α1 loop) and helix α5—that are reflected by an increase in the α1-to-α5 distance (∼10 Å in prefusion to ∼16 Å in postfusion) and an α1–(β2–α1 loop) separation (∼10 Å to ∼14 Å) (Fig. 1a). Rather than stabilizing RR1 by brute-force tethering alone, we asked whether the prefusion trimer could be arrested by rationally reinforcing the allosteric "hinge" that licenses the transition while simultaneously shoring up the interprotomer interface that stabilizes the global head.

**Fig. 1.**
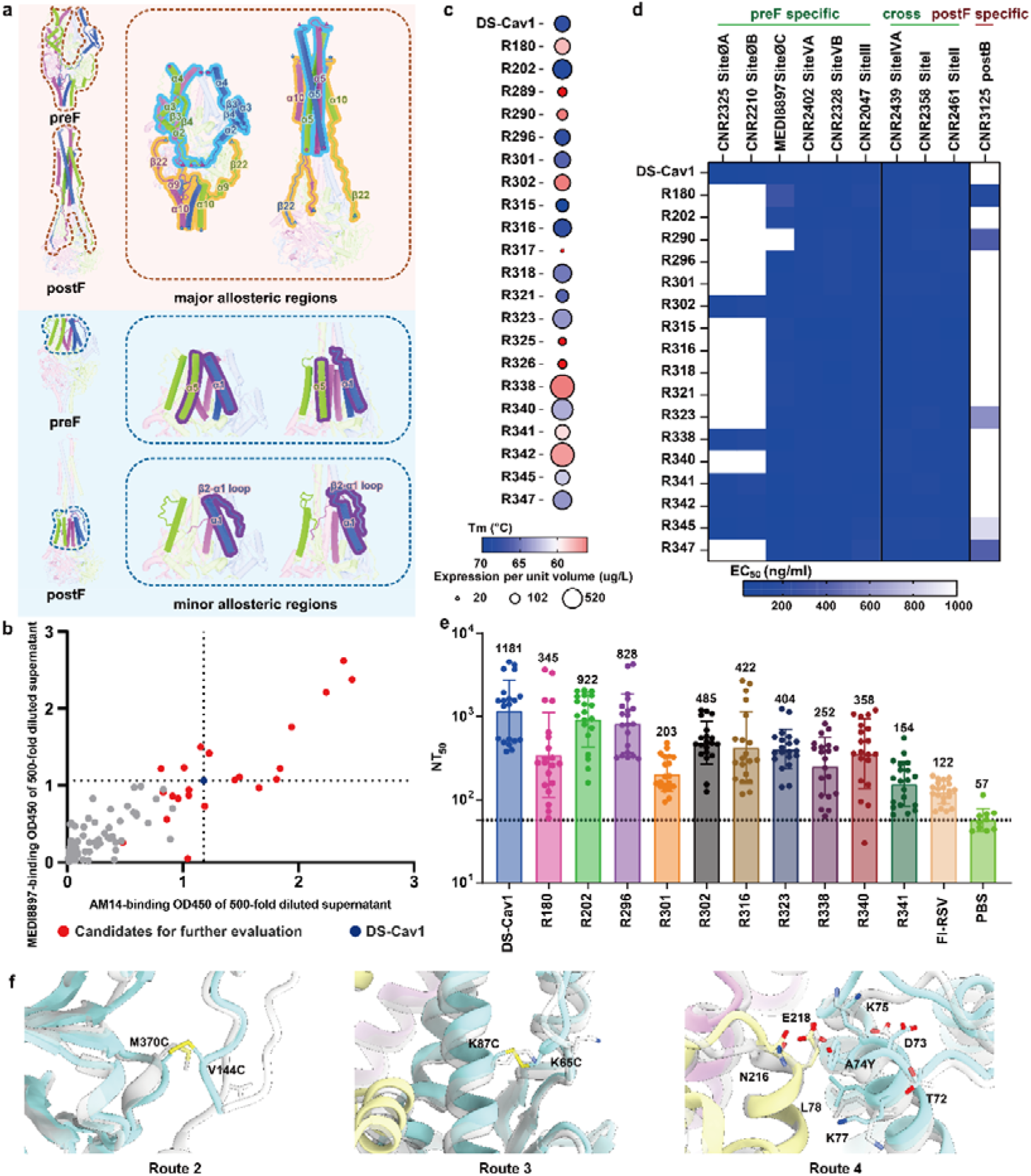
Structure-guided design and screening of prefusion RSV F immunogen candidates. **a,** Structural comparison of RSV F in the prefusion and postfusion conformations identifies conformationally dynamic regions targeted for stabilization. Large-scale refolding involves RR1 collapse into the postfusion α5 helix and RR2-mediated six-helix bundle closure, while local breathing motions around α1, the β2–α1 loop, and α5 defined major and minor allosteric regions for prefusion-stabilizing design. **b**, Primary antigenicity-based screening of designed variants by binding to AM14 and MEDI8897 using 500-fold diluted culture supernatants. Red dots indicate candidates selected for further evaluation, and DS-Cav1 is shown as a reference control. **c,** Bubble plot of expression yield and thermal stability of selected candidates. Circle size represents expression per unit culture volume, and color indicates melting temperature. **d,** Heatmap of antibody-binding EC_50_ values showing the antigenic profiles of selected candidates against prefusion-specific, pre/postfusion cross-reactive, and postfusion-specific monoclonal antibodies. **e,** Serum neutralizing antibody titers elicited by selected candidates in BALB/c after two immunizations. Neutralizing titers are shown as NT_50_ values, with DS-Cav1, FI-RSV, and PBS included as controls. Bars indicate geometric mean titers with error bars representing dispersion among animals (n = 20, n = 10 for PBS control). **f,** Cartoon representation of R296 and DS-Cav1 highlighting local structural changes around the R296 stabilizing mutations. Enlarged views show the engineered disulfide bonds and local packing interactions introduced in R296, with DS-Cav1 shown as a prefusion RSV F reference.

We implemented a four-step screen to convert structure-guided hypotheses into lead antigens (Extended Data Fig. 1): **(i)** *In silico*prioritization. ThermoNet (structure-based ΔΔ*G* prediction) ranked tolerated mutations and identified energetically stabilizing edits, whereas Rosetta scans were used to enumerate geometrically viable disulfide bridges across candidate Cα–Cα windows (e.g., α1↔α5, FP-proximal loops ↔ internal cavity walls, and trimer interface). **(ii)** Expression-level first-pass. All designs were cloned into HEK293F-compatible vectors; supernatants were screened by ELISA with two conformation-dependent probes: the trimer-specific & prefusion-sensitive antibody AM14^36,37^, and the siteØ-specific antibody MEDI8897 (nirsevimab parent) ^38^. **(iii)** Biochemical triage. Selected hits were purified and evaluated by Thermofluor (DSF) for thermal stability, then profiled with a multi-epitope antibody panel (12 functional sites) ^30^ to ensure the prefusion antigenic surface, especially siteØ, remained intact. **(iv)** *In vivo* validation. Immunogenicity (mouse), protective-efficacy surrogates, and safety/biophysical consistency checks were performed on lead candidates, with DS-Cav1^15^ and, where indicated, formalin-inactivated RSV (FI-RSV) as benchmarks. Together, this workflow couple energy-aware computation with a conformation-sensitive ELISA funnel to eliminate mis-folded or de-stabilized variants early, thereby reserving deep characterization for constructs that preserve a trimerized, prefusion and antigenically corrected configuration.

### Energy-guided mutation design across four structural strategies yields 21 validated prefusion-stabilized candidates

Guided by the ΔΔ*G*/geometry analysis, we defined four structural strategies targeting the motions that seed the prefusion-to-postfusion transition: 1), pin the postfusion C-terminal helix α10/6HB latch to resist helix dissociation; 2), stiffen the fusion peptide (FP) neighborhood and its downstream elements (α2–α4 / β3–β4) to oppose the RR1 collapse; 3), reinforce the α1–(β2–α1 loop) anchorage that gates α4/apex mobility; 4), strengthen the α5–α1 contact across protomers to raise the barrier for trimer “breathing”. For routes 1–4 we designed, respectively, 76, 91, 47, and 9 single-site or disulfide-bridge candidates; favorable edits were then rationally combined into 326 total constructs (Extended Data Table 1). Across all designs, we incorporated two established RSV prefusion F engineering features: replacement of the p27 segment with a glycine-serine (GS) linker to generate a contiguous single-chain F2–F1 antigen, following the SC-TM design strategy^16^, and fusion of a C-terminal T4 fibritin foldon domain through a flexible Gly/Ser tether to promote trimerization, following the DS-Cav1 scaffold design^15^. All constructs were expressed via a HEK293 mammalian system; supernatants were serially diluted and tested by ELISA against AM14 and MEDI8897. Positive signals clustered into four design routes and delivered tractable single-site leads, including (but not limited to), route 1: K508C, S509C, L512C, L513C, N515C/N515D, V516C, K520R; route 2: G143C, V144C, N183D, M370C, S404L/S404V, V406C; route 3: K65C, N67A/N67V/N67L, I79L, K80W/K80V, K87L/K87V/K87C, V207I; route 4: A74W, A74Y (Extended Data Fig. 1 and Extended Data Table 1). From these, 21 combinatorial mutants were assembled; ELISA confirmed that all 21 retained AM14/MEDI8897 signals not inferior to DS-Cav1 (Fig. 1b), validating that the four structural strategies can be mixed without collapsing prefusion integrity. In short, structure-guided routing of mutations—rather than undirected scan-and-pray—produces a manageable, conformation-positive combinatorial set from which a lead can be selected by biochemical and structural triage.

### Biochemical and antigenic triage narrows the pool to lead R296 with a prefusion fold confirmed by cryo-EM

The 21 candidates were purified; expression yield, Thermofluor *T*_m_, and a 10-bin epitope-panel (6 preF-specific, 3 conformation-cross-reactive, 1 postF-specific) were used to eliminate unstable or configuration-compromised constructs. Constructs R289, R317, R325, and R326 were excluded from further characterization due to low yield/poor thermal stability (Fig. 1c and Extended Data Fig. 2). ELISA-based antigenic conformation monitoring showed that several candidates, such as R180, R290, R323, R345, R347, began to adopt postfusion configuration, as revealed by strong binding to CNR3125 ^39^ (Fig. 1d and Extended Data Fig. 3). Although a number of constructs from route 3 (with mutations at positions 65, 67, 80 and 87 in the apical region) altered the local antigenicity at sites ØA/B, the proportion of site ØA antibodies induced in naturally RSV-infected individuals is extremely low (<1%), while the neutralizing activity of site ØB antibodies is relatively weak^30^, indicating that these substitutions may not affect their immunogenicity. To further verify this, we conducted an immunogenicity assessment of these constructs. Mice immunized twice (day 0/21, 6 μg, Al(OH)[adjuvant) confirmed that most candidates outperformed FI-RSV ≥ 2-fold, while R202 and R296 approached DS-Cav1 serum neutralizing titers (Fig. 1e). Because R202 exhibited a small proportion of irregular morphologies under cryo-electron microscopy (cryo-EM), we prioritized R296 for cryo-EM structure determination. The 3.31 Å map resolves the engineered mutations cleanly: the K65C–K87C disulfide locks the β2–α1 loop to α1; V144C–M370C tethers the FP-proximal region inside the trimeric cavity; and A74Y builds an interprotomer contact net (T72/D73/K75/K77/L78 with E218/N216) that further steadies the α5–α1 interprotomer hinge (Fig. 1f and Extended Data Table 2). The GS F1–F2 linker is invisible (flexible), and the Cα RMSD over 353 residues versus PDB 4MMU (DS-Cav1) is 0.817 Å, confirming the overall prefusion fold is retained without distortion (Fig. 1f and Extended Data Fig. 4). Confirmed by expression, stability, antigenic and immunogenic evaluations as well as structural verification, R296 was identified as a lead candidate, making it suitable for downstream mRNA-format or protein-format vaccine development.

### The R296 mRNA vaccine elicits potent, durable, and broadly protective immunity across preclinical models

We first evaluated the immunogenicity and protective efficacy of an LNP-encapsulated mRNA vaccine encoding the stabilized prefusion RSV F immunogen R296 in BALB/c mice, using the commercial vaccine Arexvy (GSK, 12 μg antigen) as a prefusion benchmark and FI-RSV as a non-prefusion control (Fig. 2a). On day 35, post-immunization neutralizing titers (NT[[) exceeded 10,000 in all mRNA groups except the 1 μg cohort, with the 5-, 10-, and 20-μg dose arms reaching 17,798–18,396, which were slightly higher than that of the Arexvy comparator (14,533), although the difference was not statistically significant (Fig. 2b). Cellular immunity profiling revealed a Th1-biased response: compared to Arexvy, splenic IFN-γ and IL-2 production were elevated approximately 4-, 7-, and 8-fold in the 5-, 10-, and 20-μg groups, respectively, while IL-4 increased only ∼2-fold at doses ≥ 10 μg, remaining within a non-Th2–dominant range (Fig. 2c). To assess protection, mice were challenged intranasally with RSV A2 on day 49, and lungs were harvested on day 54. No detectable viral load was monitored in any vaccinated group (limit of detection, LoD), whereas PBS controls remained ∼10^4^ titers (Fig. 2d). Thus, mRNA-encoded R296 elicited robust, dose-dependent neutralizing titers and Th1-skewed cellular immunity, achieving clearance of detectable infectious virus in murine lungs at lower doses.

**Fig. 2.**
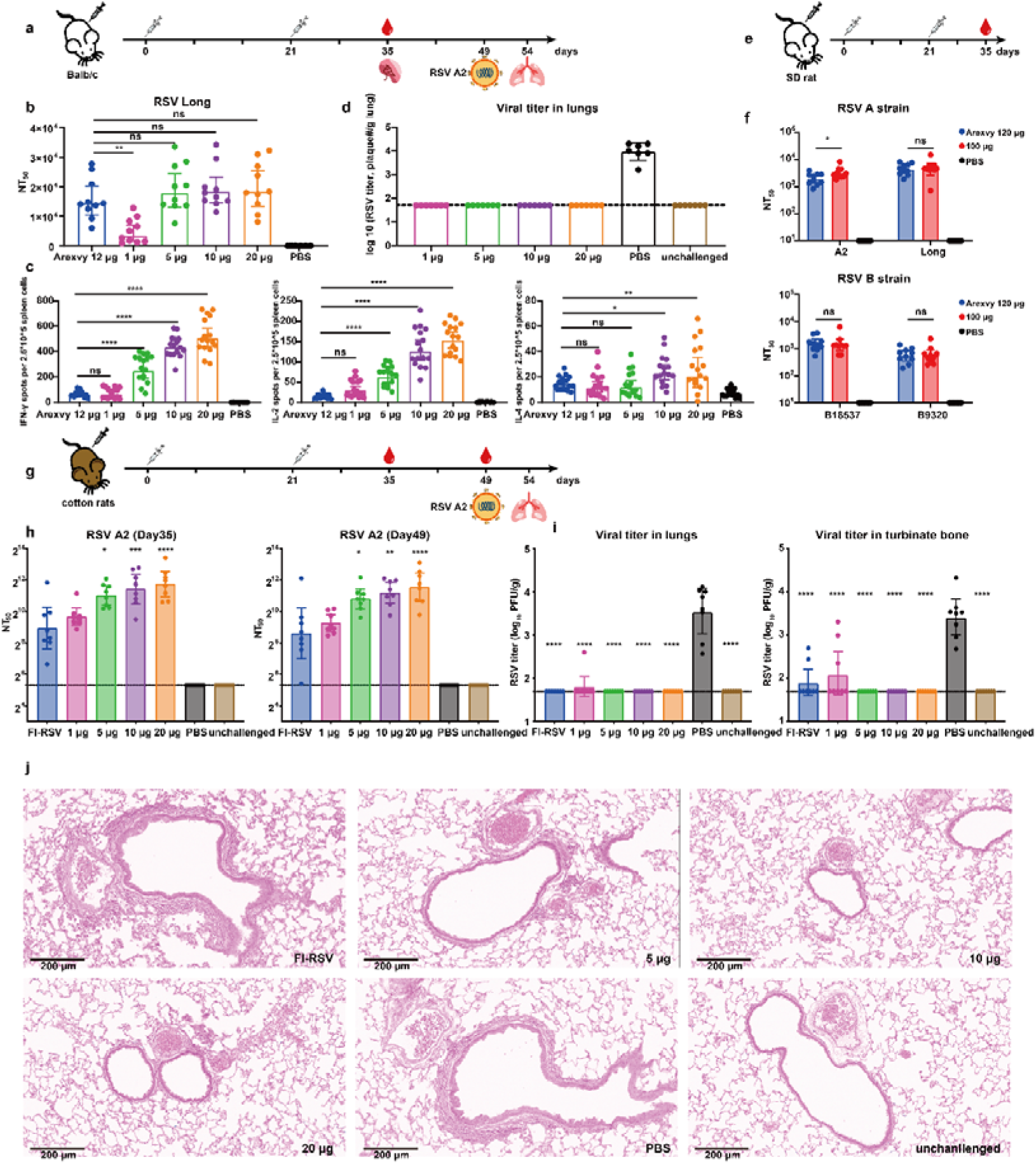
Protective efficacy of R296 mRNA vaccination in multiple animal models. **a,** Schematic of the BALB/c mouse immunization, serum collection, RSV A2 challenge, and tissue collection schedule. **b,** Serum neutralizing antibody titers against RSV Long after two immunizations with different doses of R296 mRNA (n = 10). Arexvy and PBS were included as positive and negative controls, respectively. **c,** RSV F-specific cellular immune responses measured by ELISpot. IFN-γ-, IL-2-, and IL-4-secreting cells are shown for each immunization group (n = 16). **d,** Lung viral loads in BALB/c challenged with RSV A2 after two immunizations with R296 mRNA. Viral titers were measured in lung tissues after challenge, with PBS and unchallenged mice included as controls (n = 7). **e,** Schematic of the SD rat immunization and serum collection schedule. Rats were immunized twice on days 0 and 21, and sera were collected on day 35. **f,** Serum neutralizing antibody titers in SD rats against representative RSV A and RSV B strains after immunization with R296 mRNA or Arexvy (n = 10). **g,** Schematic of the cotton rat immunization, serum collection, RSV A2 challenge, and tissue collection schedule. Cotton rats were immunized twice on days 0 and 21, challenged with RSV A2 on day 49, and lungs and nasal turbinates were collected on day 54. **h,** Serum neutralizing antibody titers against RSV A2 in cotton rats on days 35 and 49 after immunization with R296 mRNA (n = 8). FI-RSV, PBS, and unchallenged animals were included as controls. **i,** Viral titers in the lungs and nasal turbinate bones of cotton rats on day 54 after RSV A2 challenge (n = 8). **j,** Representative H&E-stained lung sections from cotton rats showing pulmonary pathology after RSV A2 challenge. R296 mRNA-vaccinated groups showed reduced inflammatory changes compared with FI-RSV and PBS controls (magnification, ×144; scale bars, 200 μm). Data are shown as individual animals with bars indicating group means and error bars indicating dispersion. Dotted lines indicate the lower limit of detection. Statistical significance is indicated as ns, not significant; *P < 0.05; **P < 0.01; ***P < 0.001; ****P < 0.0001.

Given the importance of antigenic diversity, we assessed cross-neutralization breadth and durability in Sprague-Dawley rats using a 0/21-day regimen (100 μg/dose), with Arexvy (120 μg antigen) as a comparator (Fig. 2e). Day-35 sera demonstrated potent neutralization against subgroup A strains, with NT[[values exceeding 3,215 and 4,487 against A2 and Long, respectively, both superior to the comparator (Fig. 2f). Against subgroup B strains (B18537 and B9320), NT[[values declined to 1,371 and 567, respectively, reflecting the A2-homologous sequence backbone; however, these titers remained comparable to the comparator (Fig. 2f). Longitudinal durability assessments via serial bleeds at predefined time points (days 35, 49, 63, 91, and 119) demonstrated that the induced neutralizing titers remained persistently high, with a reduction of less than 50% over the ∼4-month observation period (Extended Data Fig. 5a).

Following initial rodent validation, we further evaluated R296 mRNA performance in cotton rats (*Sigmodon hispidus*), a physiologically relevant model of human RSV infection (Fig. 2g). Animals received intramuscular prime-boost immunization (days 0 and 21) with FI-RSV or 1, 5, 10, or 20 μg of R296 mRNA. PreF-specific binding antibody titers peaked on day 35 and persisted through day 49, with titers ∼10-fold (1 μg) to ∼20-fold (5–20 μg) above FI-RSV controls (Extended Data Fig. 5b). Neutralizing titers rose dose-dependently, reaching NT[[values of 810 (1 μg), 2,087 (5 μg), 2,717 (10 μg), and 3,422 (20 μg) by day 35 (Fig. 2h). Following intranasal RSV A2 challenge (1 × 10[PFU) on day 49, lung and nasal turbinate tissues were harvested on day 54. In lungs, plaque assays revealed a ∼1.75 log[[PFU/g reduction in the 1 μg R296 group relative to PBS, with viral RNA detected in only one animal; all higher-dose groups showed complete clearance of detectable infectious virus (Fig. 2i). Similarly, in nasal turbinates, the 1 μg dose reduced viral titers by ∼1.28 log[[PFU/g, whereas doses ≥ 5 μg resulted in complete clearance of detectable infectious virus (Fig. 2i). Histopathological analysis of perivascular, peribronchial, alveolar, and interstitial compartments showed exacerbated inflammation in FI-RSV controls, whereas all R296 mRNA groups—particularly those receiving ≥ 5 μg—exhibited markedly reduced pathology, approaching a minimal scattered inflammatory pattern (Fig. 2j and Extended Data Fig. 5c). Together, these data demonstrate that R296 mRNA elicits dose-dependent systemic neutralization that translates into robust reductions in both lower and upper respiratory tract viral loads, with histopathology consistent with protection rather than VAERD.

### Safety assessment confirms absence of Th2-biased pathology and a favorable biodistribution profile

We next characterized the safety profile of R296 mRNA *in vivo*. Cytokine profiling revealed no evidence of a VAERD signature: whereas FI-RSV induced canonical Th2 polarization (elevated IL-4, IL-5, and IL-13) in lung tissue, all mRNA dose groups lacked this response and instead restored IFN-γ expression (Fig. 3a). Lung histopathology paralleled these findings, with FI-RSV exacerbating total histopathology score, whereas mRNA vaccination reduced total inflammation in a dose-dependent manner, with doses ≥ 5 μg improving peribronchiolar and interstitial pathology (Fig. 3b). To evaluate systemic exposure, we assessed biodistribution and metabolic clearance in SD rats following a single 50-μg intramuscular dose. mRNA transcripts peaked in tissues and whole blood within 2–24 hours and underwent rapid washout. The injection-site muscle represented the major early depot, whereas by day 21, quantifiable mRNA was largely restricted to immune-draining sites, including inguinal lymph nodes, spleen, and mesenteric lymph nodes. Brain, contralateral muscle, gonads, and thymus remained at or below the limit of quantification (LoQ) (Fig. 3, c and d). The ionizable lipid SM-102 exhibited broad early tissue distribution but concentrated in lymph nodes, spleen, and injection muscle (∼9–305× plasma levels), with low-to-moderate liver and adrenal spillover and negligible gonadal signal, clearing to low or undetectable levels by day 21 (Fig. 3, c and d). The helper lipid mPEG-DMG-2K remained low throughout, with quantifiable signal restricted to plasma, injection-site muscle, and inguinal lymph nodes (Fig. 3c, right). Finally, lyophilization stability testing demonstrated that post-lyophilization immunogenicity was statistically indistinguishable from pre-lyophilization baselines across 5- and 10-μg dose tiers (Extended Data Fig. 5d). Together, these data indicate that R296 mRNA shows no evidence of Th2/VAERD-associated pathology under the tested conditions, exhibits a self-limiting, muscle-centric biodistribution profile without persistent off-target deposition, and supports a thermostable dry formulation compatible with global deployment.

**Fig. 3.**
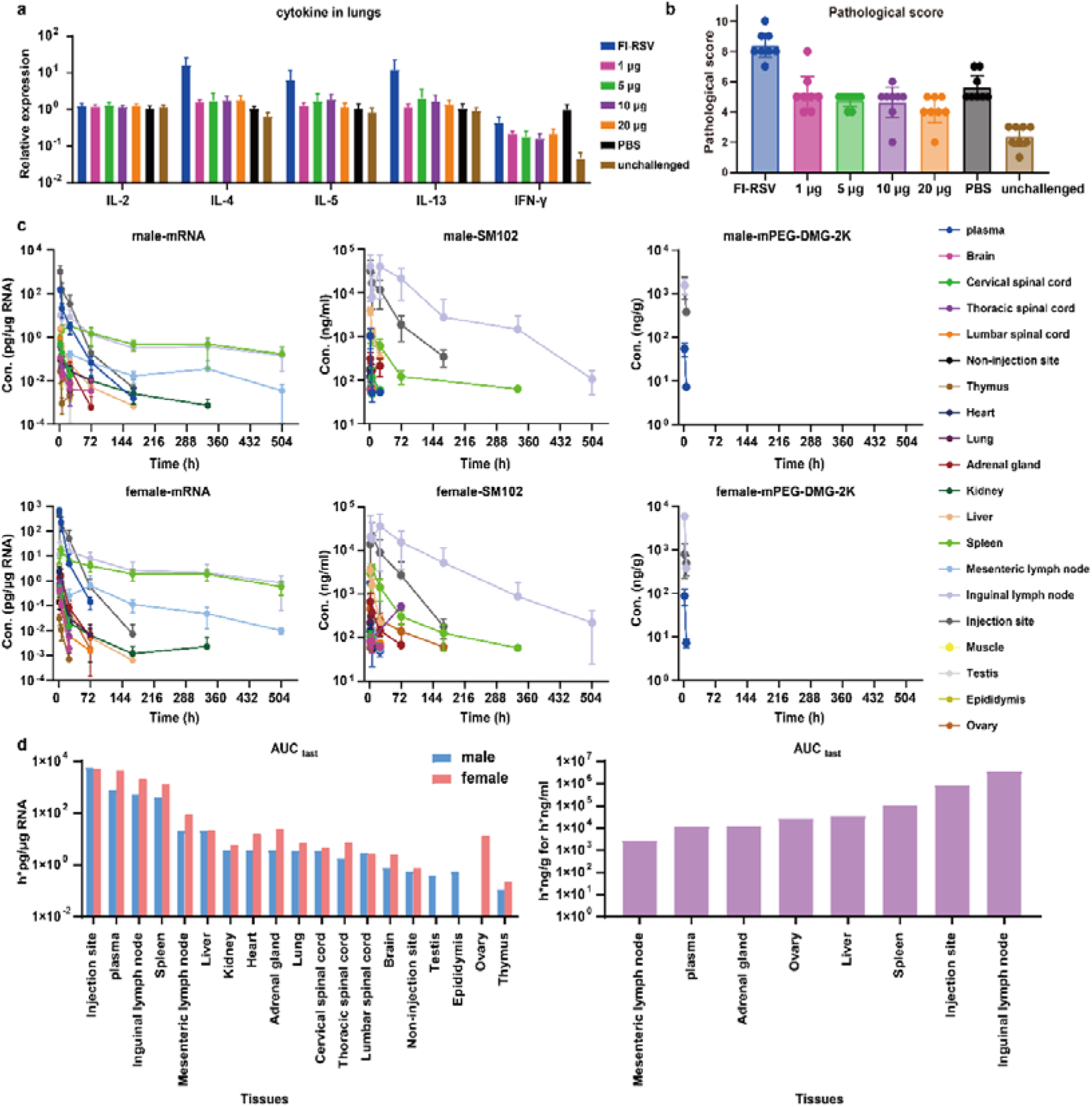
Safety assessment and pharmacokinetic profiling of R296 mRNA vaccination. **a,** Lung cytokine expression after RSV A2 challenge in cotton rats immunized with R296 mRNA or control formulations. Relative expression levels of Th1- and Th2-associated cytokines are shown to evaluate vaccine-associated enhanced respiratory disease-related inflammatory responses (n = 8). **b,** Semi-quantitative pathological scoring of lung tissues after RSV A2 challenge. R296 mRNA vaccination reduced lung pathology compared with FI-RSV and PBS controls, with unchallenged animals included as the baseline control (n = 8). **c,** Tissue distribution and clearance kinetics of mRNA, SM-102, and mPEG-DMG-2K in male and female SD rats after intramuscular administration of R296 mRNA. Concentrations were measured in plasma and major tissues over time (n = 5). **d,** Tissue exposure of mRNA and SM-102 calculated as AUC_last_ in male and female SD rats. Bars show the cumulative exposure of mRNA and SM-102 across indicated tissues. Data are shown as mean ± SEM where applicable (n = 5).

### Epitope-guided stalkless nanoparticle immunogens confer protective efficacy without Th2/VAERD-type pathology

Recent clinical holds on pediatric RSV mRNA programs, prompted by a numerical imbalance in severe lower respiratory tract illness among 5- to 8-month-old infants, highlight the need to consider “antigenic purity” as a critical safety determinant extending beyond global prefusion stability to the specific epitopes presented^12^. Leveraging our previously defined 12-functional-class antibody landscape (Fig. 4a), we mapped neutralizing potency onto the preF structure and observed a concentration of high-activity clones targeting the apical head (sites ØA–ØC/V), whereas the membrane-proximal stalk (sites IV/I/VI) predominantly harbored low-potency or non-neutralizing antibodies (Fig. 4, b and c). Correlating with this, approximately 80% of toddler antibodies target central-to-basal, low-potency epitopes (III/IV/I/VI) engaged by germline-encoded clones with minimal somatic hypermutation, while adult repertoires concentrate on apical, high-potency sites (Ø/V), where top-tier ØA/ØC neutralizers constitute ∼24% of clones yet remain nearly undetectable in toddlers (< 1%)—a parsimonious and plausible mechanistic explanation for historic pediatric vaccine failures^30^. To enrich for protective epitopes, we designed truncation constructs that excise the stalk while retaining a contiguous head-like unit anchored on antigenic site III (Fig. 4d). To compensate for stability loss from truncation, we generated 42 stabilizing variants incorporating point mutations and micro-insertions (Extended Data Table 3). Although constructs Head11 and Head38 were the top expressers (Extended Data Fig. 6a), the ∼1/6 reduction in molecular mass was predicted to have intrinsically low B-cell avidity. We therefore displayed construct Head38 on the I53-50AB nanoparticle^40^ via variable-length Gly/Ser linkers (Fig. 4e). Screening four linker lengths (6, 10, 14, 18 aa) confirmed that the 14-aa candidate (Head38-50AB-3) retained high-affinity recognition for head-restricted, preF-potent epitopes and adopted the expected head-like fold with preserved siteØ/V/II topology (Extended Data Fig. 6, b and c)^39^. To further validate its structural integrity and epitope presentation, we determined a 4.4 Å cryo-EM structure of Head38 in complex with CNR2292 and motavizumab^39,41^. The structure confirmed that the stalk-excised immunogen maintained the intended head-like architecture and preserved the conformational accessibility of high-potency neutralizing epitopes (Fig. 4f, Extended Data Fig. 7 and Extended Data Table 2). This epitope-guided approach yielded an immunogen enriched for high-potency apex epitopes and depleted of stalk neighborhoods historically associated with non-protective or VAERD-prone immunity.

**Fig. 4.**
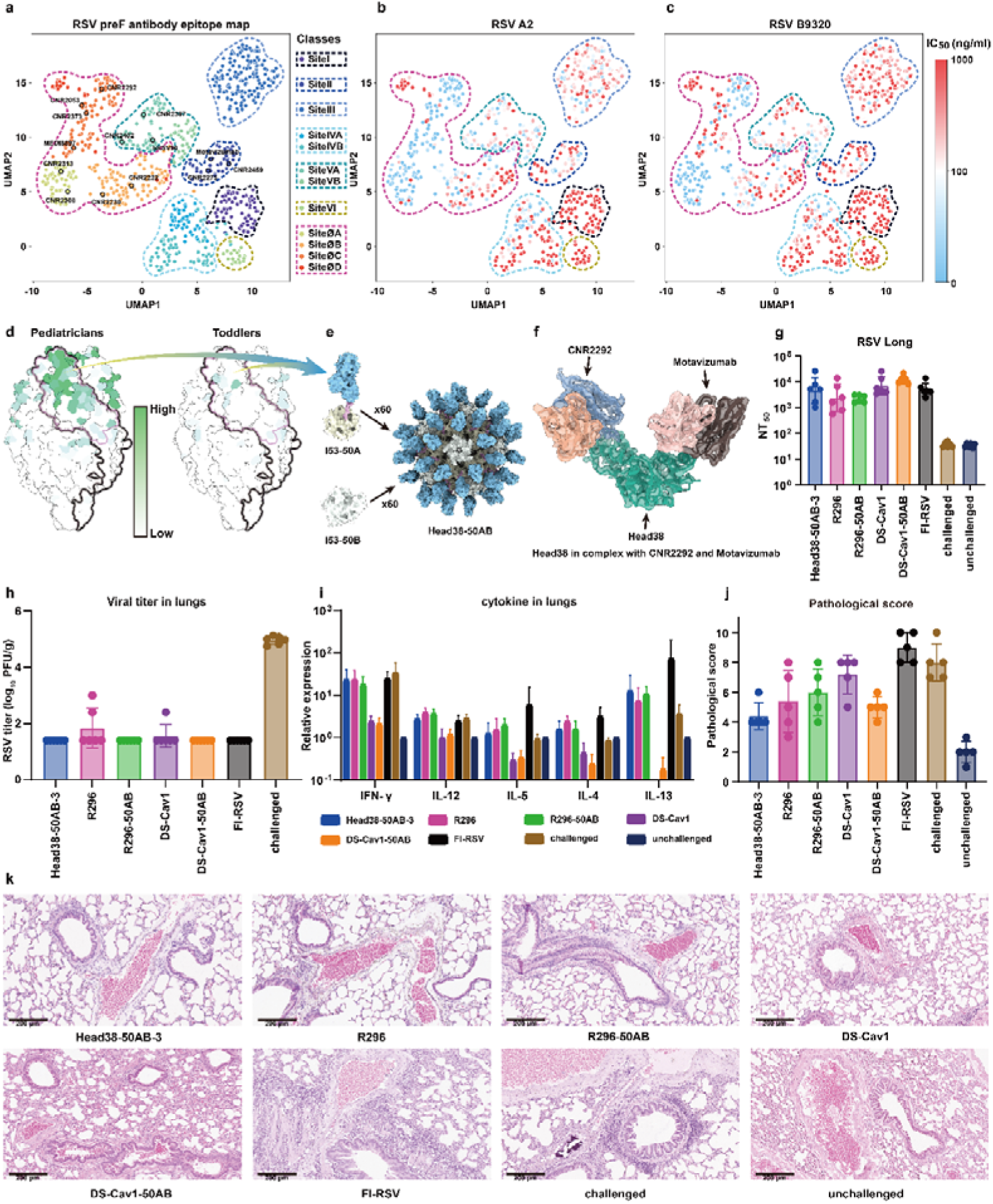
Design, structural validation, and protective efficacy of truncated RSV F Head38 displayed on I53-50AB nanoparticles. **a-d,** Analyses based on previously published RSV F–specific antibody datasets ^1^, replotted here to provide the rationale for epitope-focused immunogen design. **a,** UMAP projection showing the classification of RSV F-specific antibodies based on epitope-targeting patterns and functional features. **b,c,** UMAPs colored by neutralizing activity against RSV A2 **(b)** and RSV B9320 **(c)**, revealing the distribution of neutralizing potency across antibody clusters. **d,** RSV F surface representations colored by the neutralizing activity of antibodies targeting different antigenic regions in pediatricians and toddlers. Strongly neutralizing responses are enriched on the prefusion apex in pediatricians, whereas toddler-derived antibodies show weaker and more dispersed neutralizing activity. **e,** Schematic of the truncation strategy used to generate head-focused RSV F immunogens and their multivalent display on the I53-50AB nanoparticle through flexible Gly/Ser linkers. **f,** Cryo-EM structure of the Head38 immunogen in complex with CNR2292 and motavizumab. **g-k,** Immunogenicity, protective efficacy, inflammatory responses, and lung pathology in cotton rats immunized with truncated RSV F nanoparticle candidates or control immunogens and challenged with RSV A2. **(g)** Serum neutralizing antibody titers (n = 6). **(h)** Lung viral titers after RSV A2 challenge (n = 6). **(i)** Lung cytokine expression after challenge, assessing Th1/Th2-associated inflammatory responses and vaccine-associated enhanced respiratory disease-related signatures (n = 6). **(j)** Semi-quantitative pathological scoring of lung tissues (n = 5). **(k)** Representative H&E-stained lung sections after challenge (magnification, ×144; scale bars, 200 μm). Data are shown as individual animals with bars indicating group means and error bars indicating dispersion where applicable. Dotted lines indicate the lower limit of detection where shown. Statistical significance is indicated as ns, not significant; *P < 0.05; **P < 0.01; ***P < 0.001; ****P < 0.0001.

We evaluated the lead candidate, Head38-50AB-3, alongside comparator immunogens—including soluble R296, R296-50AB, DS-Cav1-50AB, and DS-Cav1 protein—in cotton rats (6 μg/dose, days 0/21; RSV A2 challenge on day 49; harvest on day 54). Head38-50AB-3 elicited neutralizing titers statistically on par with full-length preF trimer and nanoparticle benchmarks (Fig. 4g). Upon challenge, all vaccine arms reduced lung viral load by 3.22–3.56 log[[PFU/g compared to PBS controls, with several groups showing complete viral clearance of detectable infectious virus (Fig. 4h). Crucially, qPCR analysis revealed no significant elevation of Th2 cytokines (IL-4, IL-5, IL-13) across any vaccine group (Fig. 4i). Histopathological examination showed no evidence of enhanced peribronchiolar or interstitial inflammation attributable to vaccination, whereas FI-RSV controls remained the sole group exhibiting the classical VAERD-type pattern (Fig. 4, j and k). Together, these data demonstrate that epitope-guided stalk excision and nanoparticle display eliminate exposure to stalk epitopes associated with historic VAERD risk while inducing potent, protective neutralization.

### A ciliary-adhesive LNP enables intranasal RSV mRNA vaccination with systemic and mucosal immunity

Systemic immunization with prefusion F vaccines elicits robust serum neutralizing antibodies but limited respiratory mucosal immunity, resulting in weak sIgA production and insufficient tissue-resident memory responses^22,31,42^. To overcome this limitation, we engineered a nasal-delivery LNP (nLNP) designed to prolong ciliary adherence and promote local mRNA expression. Starting from a canonical four-component LNP architecture, we optimized ionizable-lipid candidates and lipid ratios^43,44^. Although all lead candidates met physicochemical criteria (> 90% encapsulation, ∼80–110 nm in diameter and low polydispersity), their functional performance diverged. Using intranasally delivered luciferase mRNA as a reporter, the lead formulation A3-nLNP elicited ∼11-fold higher nasal bioluminescence than SM-102–based reference nLNP at 6 hours post-dose (Fig. 5a and Extended Data Fig. S8). This reference formulation was selected because SM-102–based LNPs (SM102-nLNP) have been used in recent studies for intranasal delivery of COVID-19 and influenza mRNA antigens^45,46^, establishing A3-nLNP as the lead candidate. Cryo–transmission electron microscopy demonstrated that A3-nLNP formed uniform, spherical particles with a hydrodynamic diameter of 94.2 ± 2.4 nm; polydispersity index 0.11 ± 0.01, and > 95% mRNA encapsulation (Fig. 5b). Scanning electron microscopy of nasal tissue collected 30 minutes after dosing showed that A3-nLNP adhered along the ciliary brush from tip to base, whereas SM102-nLNP was retained within the mobile mucus layer (Fig. 5c). Consistently, kinetic profiling showed that A3-nLNP sustained high-level luciferase mRNA expression for up to 15 hours, whereas SM102-nLNP expression decayed rapidly (Fig. 5d). These data indicate that the ciliary adhesion of A3-nLNP prolongs mucosal residence and enhances local mRNA expression, supporting its potential for intranasal vaccination.

**Fig. 5.**
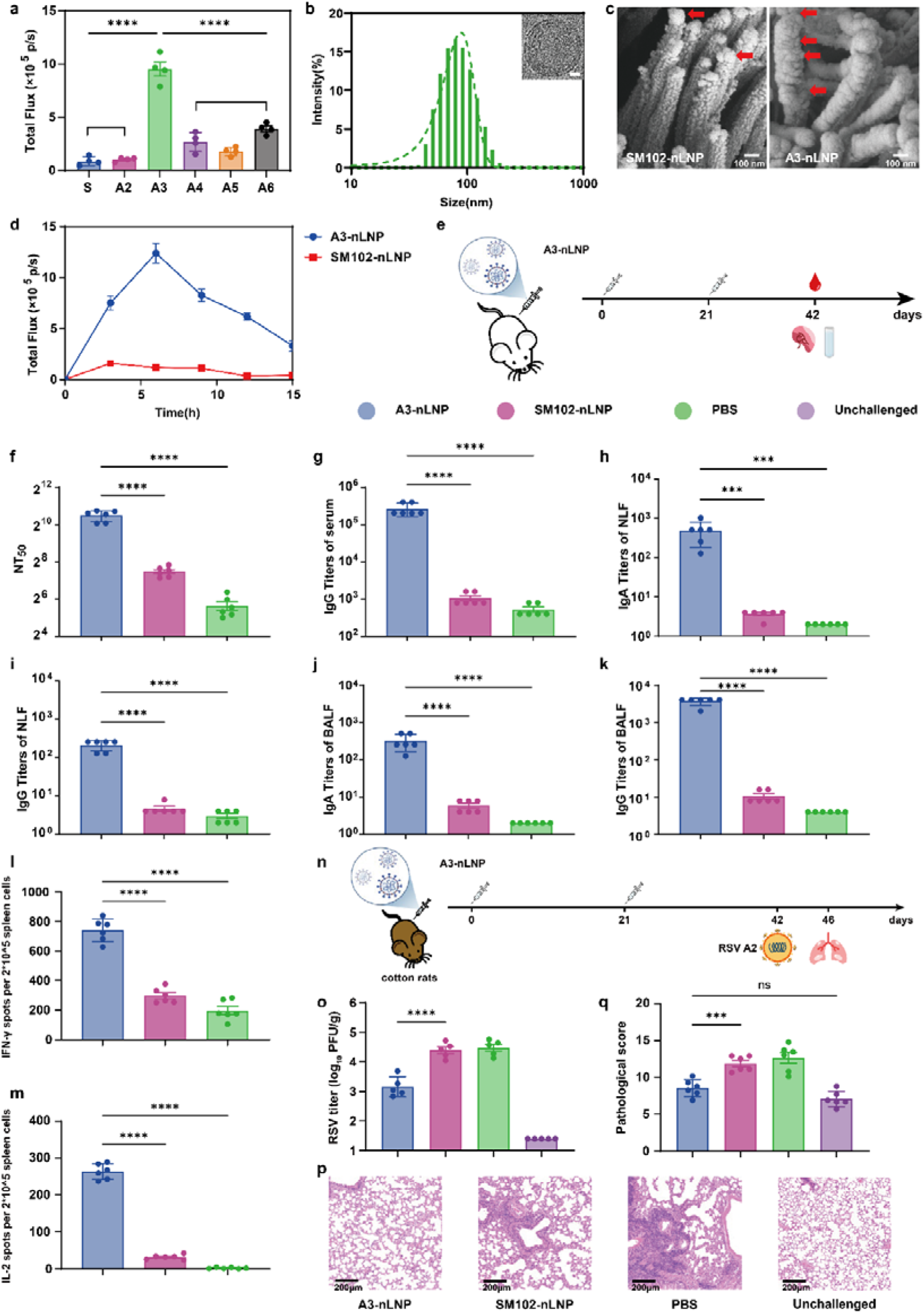
Mucosal delivery of mRNA by intranasal LNP elicits potent systemic and mucosal immunity. **a,** Total luciferase mRNA expression in the nasal cavity 6 h after intranasal administration of different LNP formulations carrying Luc mRNA. **b,** Physicochemical characterization of A3-nLNP, including hydrodynamic diameter measured by dynamic light scattering and representative cryo-TEM morphology. Scale bar, 10 nm. **c,** Representative SEM images showing the attachment of A3-nLNP or SM102-nLNP to nasal cilia 30 min after intranasal administration. Red arrows indicate LNP particles associated with nasal cilia. Scale bars, 100 nm. **d,** Kinetics of nasal luciferase expression after intranasal delivery of A3-nLNP or SM102-nLNP. **e,** Schematic of the mouse intranasal immunization and sample collection schedule. BALB/c mice were immunized twice on days 0 and 21 with R296 mRNA formulated in LNP, and serum, nasal lavage fluid, bronchoalveolar lavage fluid, and spleens were collected for immune analyses. **f,** Serum neutralizing antibody titers (NT[[) against RSV A2 pseudovirus after immunization (n = 6). **g-k,** RSV F-specific antibody responses measured by ELISA, including serum IgG (n = 6) **(g)**, nasal lavage fluid IgA **(h)**, nasal lavage fluid IgG **(i)**, bronchoalveolar lavage fluid IgA **(j)**, and bronchoalveolar lavage fluid IgG (K). **l,m,** ELISpot analysis of antigen-specific IFN-γ- producing **(l)** and IL-2-producing (n = 6) **(m)** splenocytes after peptide restimulation. **n,** Schematic of the cotton rat immunization and RSV challenge experiment. Cotton rats were immunized twice on days 0 and 21, challenged intranasally with RSV on day 42, and lung tissues were collected 4 days after challenge. **o,** Lung viral loads in cotton rats 4 days after RSV challenge, expressed as log[[PFU/g lung tissue (n = 5). **p,** Representative H&E-stained lung sections showing pulmonary pathology after RSV challenge. Scale bars, 200 μm. Data are shown as mean ± SEM. **q,**Semi-quantitative histopathological scoring of lung sections after RSV challenge (n = 6). Statistical significance was determined by one-way ANOVA with Tukey’s multiple-comparison test. ns, not significant; *P < 0.05; **P < 0.01; ***P < 0.001; ****P < 0.0001.

We next encapsulated R296 mRNA into A3-nLNP and immunized BALB/c mice intranasally on days 0 and 21 at a dose of 10 μg mRNA, with CpG adjuvant. Intranasal A3-nLNP elicited robust serum neutralizing titers (NT[[), compared with SM102-nLNP and PBS controls (Fig. 5f), while also inducing robust systemic anti-preF IgG responses (Fig. 5g). In the mucosal compartment, nasal lavage fluid (NLF) showed an ∼100-fold increase in sIgA and a ∼45-fold increase in anti-preF-specific IgG, while bronchoalveolar lavage fluid (BALF) showed a ∼53-fold increase in sIgA (Fig. 5h–k). These results demonstrate effective engagement of both nasal- and bronchus-associated mucosal antibody barriers at the respiratory portal of entry. Splenic ELISpot suggested a Th1-polarized systemic cellular response, with significantly elevated frequencies of IFN-γ- and IL-2-producing splenocytes (Fig. 5, l and m and Extended Data Fig. 9). Cytokine analysis of culture supernatants showed elevated IFN-γ, IL-2, TNF-α, and IL-17A levels, supporting a type 1-biased immune profile (Extended Data Fig. 10).

In cotton rats, prime-boost immunization with A3-nLNP (10 μg, days 0/21) (Fig. 5n) reduced lung viral titers following RSV A2 challenge at week 6, compared with SM102-nLNP (Fig. 5o). Histopathological analysis showed that the PBS and SM102-nLNP groups retained diffuse perivascular and peribronchiolar infiltrates with alveolar edema, whereas the A3-nLNP group showed preserved alveolar architecture (Fig. 5p). Semi-quantitative pathology scoring confirmed a significant reduction in total inflammation for A3-nLNP relative to controls (Fig. 5q). Together, these data demonstrate that ciliary-adhesive intranasal delivery of R296 mRNA using A3-nLNP induces potent mucosal sIgA and IgG responses at the respiratory portal of entry, while also eliciting systemic neutralizing antibodies and IFN-γ-associated cellular immunity.

## DISCUSSION

This study describes a vertically integrated, structure-guided pipeline that advances RSV vaccine design beyond empirical “stabilize-and-hope” approaches toward a deterministic, risk-aware engineering framework. Central to this workflow is a four-pronged computational-to-biochemical screening funnel anchored by allosteric mapping of prefusion F dynamics—specifically α1–α5 hinge motions and RR1/RR2 coupling—and translated through ThermoNet ΔΔG prioritization, Rosetta-enabled disulfide and interprotomer contact design, and conformation-sensitive ELISA triage using prefusion- and trimer-specific probes (e.g., AM14, MEDI8897) alongside a multi-epitope antibody panel (Fig. 1). By treating antigenicity and stability as co-equal readouts, this strategy front-loads attrition: misfolded, destabilized, or conformationally leaky constructs are discarded before resource-intensive in vivo validation. The result is a reusable design–measure–triage–verify loop, wherein cryo-EM validation (e.g., the K65C–K87C lock, FP-proximal tether, and A74Y network in R296) closes the feedback loop between in silico rationale and in situ structural reality.

A second focus of this work is the recognition that prefusion stabilization alone does not ensure pediatric safety. Recent clinical holds on infant RSV mRNA programs underscore the necessity of treating epitope composition as a first-order safety variable^12^. Addressing this, we previously mapped a 12-functional-class antibody landscape onto the preF topography^30^ and applied an epitope-guided stalk-excision strategy. This approach enriches apical, high-potency neutralizing surfaces while depleting membrane-proximal stalk neighborhoods that disproportionately harbor low-potency or non-neutralizing epitopes implicated in Th2-biased or putative VAERD priming^26,47^. When displayed on an I53-50AB nanoparticle (Head38-50AB-3), this head-only immunogen restored avidity and retained native site Ø/V/II topology (Fig. 4), while eliminating putative structural substrates for immunopathology. In cotton rats, Head38-50AB-3 achieved neutralizing titers and clearance of detectable infectious virus in lungs on par with full-length preF benchmarks, without the Th2 cytokine elevations or peribronchiolar/interstitial pathology characteristic of FI-RSV controls. These data support a refined paradigm: “antigenic purity” is defined not merely by the presence of a prefusion fold, but by the specific repertoire of exposed epitopes, with rational excision offering a path to pediatric-viable candidates.

A significant hurdle in RSV prophylaxis remains the generation of mucosal immunity. Systemic preF vaccines excel at generating serum neutralization but often fail to seed durable sIgA or tissue-resident memory at the primary site of RSV entry^22,31^. The A3-nLNP platform described here offers a potential strategy to mitigate this gap by enhancing intranasal mRNA delivery through ciliary adhesion (Fig. 5). Compared with the SM-102–based reference nLNP, A3-nLNP showed stronger ciliary adhesion, prolonged nasal luciferase expression, and enhanced local transfection after intranasal delivery. When loaded with R296 mRNA, A3-nLNP elicited systemic neutralizing antibodies, anti-preF IgG, mucosal sIgA in nasal and bronchoalveolar lavages, and a Th1-biased cellular response marked by increased IFN-γ and IL-2. In cotton rats, ntranasal A3-nLNP vaccination reduced lung RSV A2 titers and mitigated inflammatory lung pathology compared with control formulations, supporting the functional relevance of this mucosal delivery strategy. Nevertheless, the relationship between enhanced ciliary residence, local mRNA expression, mucosal antibody induction, and protection from upper-airway infection requires further investigation. Future studies are expected to take a deeper investigation on tissue-resident T-cell subsets in the nasal and lung mucosa, and evaluate challenge models that more closely reflect natural RSV exposure and transmission.

Several limitations warrant consideration. Although cotton rats reproduce several pathological features of human RSV infection, no small-animal model fully recapitulates infant immune ontogeny, maternal antibody interference, or the narrow developmental window associated with susceptibility to VAERD^12,48^. Additionally, while rat studies indicate stable neutralizing titers over ∼4 months and lyophilization maintains potency, true durability in target populations, particularly older adults with immunosenescence or infants with waning maternal IgG, remains unestablished. Breadth is another constraint: although the A2-homologous backbone provides strong subgroup A neutralization, real-world effectiveness depends on performance against circulating, drift-capable RSV strains, necessitating broader panel-based neutralization and escape-mapping studies. Finally, the safety envelope of the nLNP formulation, including nerve epithelium tolerance, olfactory bulb translocation risk, and chronic inflammatory potential, will require rigorous nonclinical toxicology and careful dose-escalation studies.

In summary, this work outlines a next-generation “design–conformation–safety–delivery” continuum for RSV. By integrating allosteric-hinge engineering (R296), epitope-guided stalkless presentation (Head38-50AB-3), and a ciliary-adhesive intranasal mRNA platform (A3 LNP), we offer a portfolio-style strategy pairing thermostable, curated immunogens with anatomically matched delivery vehicles. Notably, the lead candidate R296 has recently advanced into a Phase 1 clinical trial (CTR20260840), providing a crucial translational test of this structure-guided design logic in humans. If these early-stage data confirm the safety and immunogenicity profiles observed preclinically, particularly regarding infant-specific safety margins and mucosal durability, this approach could help close two persistent gaps: enabling pediatric-viable RSV antigens and achieving scalable mucosal protection without compromising the potent neutralization the field has refined over the past decade.

## Acknowledgments

We thank X. Huang, X. Li, B. Zhu and L. Chen for cryo-EM data collection at the Center for Biological imaging (CBI) in Institute of Biophysics for EM work. We also acknowledge Sinovac Life Sciences for technical assistance and valuable support throughout this work.

## Author contributions

Conceptualization: X.W., Z.L., Y.X., W.F., D.L.; Methodology: W.F., D.L., X.S., C.D., H.Z., L.J., Y.H., Z.L., Y.X., X.W.; Investigation: W.F., D.L., X.S., C.D., H.Z., L.J., Y.Z., W.M., Y.H.; Formal analysis: W.F., D.L., X.S., C.D., H.Z., L.J., Y.Z., W.M., Y.H.; Visualization: W.F., D.L., X.S., C.D.; Funding acquisition: X.W., Z.L., Y.X.; Project administration: X.W., Z.L., Y.X.; Supervision: X.W., Z.L., Y.X.; Writing – original draft: W.F., D.L., Z.L., X.W.; Writing – review & editing: W.F., D.L., X.S., C.D., H.Z., L.J., Y.Z., W.M., Y.H., Z.L., Y.X., X.W.

## References and Notes

1 Shi, T., Vennard, S., Jasiewicz, F., Brogden, R. & Nair, H. Disease Burden Estimates of Respiratory Syncytial Virus related Acute Respiratory Infections in Adults With Comorbidity: A Systematic Review and Meta-Analysis. J Infect Dis 226, S17–s21 (2022).

2 Li, Y., et al. Global, regional, and national disease burden estimates of acute lower respiratory infections due to respiratory syncytial virus in children younger than 5 years in 2019: a systematic analysis. Lancet 399, 2047–2064 (2022).

3 Scheltema, N. M., et al. Global respiratory syncytial virus-associated mortality in young children (RSV GOLD): a retrospective case series. Lancet Glob Health 5, e984–e991 (2017).

4 Papi, A., et al. Respiratory Syncytial Virus Prefusion F Protein Vaccine in Older Adults. N Engl J Med 388, 595–608 (2023).

5 Walsh, E. E., et al. Efficacy and Safety of a Bivalent RSV Prefusion F Vaccine in Older Adults. N Engl J Med 388, 1465–1477 (2023).

6 Wilson, E., et al. Efficacy and Safety of an mRNA-Based RSV PreF Vaccine in Older Adults. N Engl J Med 389, 2233–2244 (2023).

7 McLellan, J. S. Neutralizing epitopes on the respiratory syncytial virus fusion glycoprotein. Curr Opin Virol 11, 70–75 (2015).

8 Langedijk, A. C. & Bont, L. J. Respiratory syncytial virus infection and novel interventions. Nat Rev Microbiol 21, 734–749 (2023).

9 McLellan, J. S., et al. Structure of RSV fusion glycoprotein trimer bound to a prefusion-specific neutralizing antibody. Science 340, 1113–1117 (2013).

10 Ison, M. G., et al. Efficacy and Safety of Respiratory Syncytial Virus (RSV) Prefusion F Protein Vaccine (RSVPreF3 OA) in Older Adults Over 2 RSV Seasons. Clin Infect Dis 78, 1732–1744 (2024).

11 Walsh, E. E. et al. Efficacy, Immunogenicity, and Safety of the Bivalent Respiratory Syncytial Virus (RSV) Prefusion F Vaccine in Older Adults Over 2 RSV Seasons. Clin Infect Dis 81, e680–e689 (2026).

12 Mahase, E. FDA pauses all infant RSV vaccine trials after rise in severe illnesses. Bmj 387, q2852 (2024).

13 Britton, A., et al. Use of Respiratory Syncytial Virus Vaccines in Adults Aged ≥60 Years: Updated Recommendations of the Advisory Committee on Immunization Practices - United States, 2024. MMWR Morb Mortal Wkly Rep 73, 696-702 (2024).

14 Kampmann, B., et al. Bivalent Prefusion F Vaccine in Pregnancy to Prevent RSV Illness in Infants. N Engl J Med 388, 1451–1464 (2023).

15 McLellan, J. S., et al. Structure-based design of a fusion glycoprotein vaccine for respiratory syncytial virus. Science 342, 592–598 (2013).

16 Krarup, A., et al. A highly stable prefusion RSV F vaccine derived from structural analysis of the fusion mechanism. Nat Commun 6, 8143 (2015).

17 Joyce, M. G., et al. Iterative structure-based improvement of a fusion-glycoprotein vaccine against RSV. Nat Struct Mol Biol 23, 811–820 (2016).

18 Huang, Q., et al. Highly scalable prefusion-stabilized RSV F vaccine with enhanced immunogenicity and robust protection. Nat Commun 16, 7805 (2025).

19 Liang, Y., et al. Mutating a flexible region of the RSV F protein can stabilize the prefusion conformation. Science 385, 1484–1491 (2024).

20 Lee, Y. Z., et al. Rational design of uncleaved prefusion-closed trimer vaccines for human respiratory syncytial virus and metapneumovirus. Nat Commun 15, 9939 (2024).

21 Jones, H. G., et al. Alternative conformations of a major antigenic site on RSV F. PLoS Pathog 15, e1007944 (2019).

22 Zohar, T., et al. Upper and lower respiratory tract correlates of protection against respiratory syncytial virus following vaccination of nonhuman primates. Cell Host Microbe 30, 41–52.e45 (2022).

23 Drysdale, S. B., et al. Priorities for developing respiratory syncytial virus vaccines in different target populations. Sci Transl Med 12 (2020).

24 Polack, F. P., et al. A role for immune complexes in enhanced respiratory syncytial virus disease. J Exp Med 196, 859–865 (2002).

25 Delgado, M. F., et al. Lack of antibody affinity maturation due to poor Toll-like receptor stimulation leads to enhanced respiratory syncytial virus disease. Nat Med 15, 34–41 (2009).

26 Knudson, C. J., Hartwig, S. M., Meyerholz, D. K. & Varga, S. M. RSV vaccine-enhanced disease is orchestrated by the combined actions of distinct CD4 T cell subsets. PLoS Pathog 11, e1004757 (2015).

27 Ngwuta, J. O., et al. Prefusion F-specific antibodies determine the magnitude of RSV neutralizing activity in human sera. Sci Transl Med 7, 309ra162 (2015).

28 Swanson, K. A., et al. A respiratory syncytial virus (RSV) F protein nanoparticle vaccine focuses antibody responses to a conserved neutralization domain. Sci Immunol 5 (2020).

29 Gilman, M. S., et al. Rapid profiling of RSV antibody repertoires from the memory B cells of naturally infected adult donors. Sci Immunol 1 (2016).

30 Deng, J. et al. Decoding protective immunity to RSV: A functional epitope landscape prescribes next-generation pediatric vaccine design. bioRxiv (2026).

31 Mettelman, R. C., Allen, E. K. & Thomas, P. G. Mucosal immune responses to infection and vaccination in the respiratory tract. Immunity 55, 749–780 (2022).

32 Li, B., Yang, Y. T., Capra, J. A. & Gerstein, M. B. Predicting changes in protein thermodynamic stability upon point mutation with deep 3D convolutional neural networks. PLoS Comput Biol 16, e1008291 (2020).

33 Leaver-Fay, A., et al. ROSETTA3: an object-oriented software suite for the simulation and design of macromolecules. Methods Enzymol 487, 545–574 (2011).

34 González-Reyes, L., et al. Cleavage of the human respiratory syncytial virus fusion protein at two distinct sites is required for activation of membrane fusion. Proc Natl Acad Sci U S A 98, 9859–9864 (2001).

35 Swanson, K. A., et al. Structural basis for immunization with postfusion respiratory syncytial virus fusion F glycoprotein (RSV F) to elicit high neutralizing antibody titers. Proc Natl Acad Sci U S A 108, 9619–9624 (2011).

36 Gilman, M. S., et al. Characterization of a Prefusion-Specific Antibody That Recognizes a Quaternary, Cleavage-Dependent Epitope on the RSV Fusion Glycoprotein. PLoS Pathog 11, e1005035 (2015).

37 Harshbarger, W., et al. Improved epitope resolution of the prefusion trimer-specific antibody AM14 bound to the RSV F glycoprotein. MAbs 13, 1955812 (2021).

38 Zhu, Q., et al. A highly potent extended half-life antibody as a potential RSV vaccine surrogate for all infants. Sci Transl Med 9 (2017).

39 Li, Y., et al. MAAD: multidimensional antiviral antibody database. Protein Cell 17, 560–572 (2026).

40 Bale, J. B., et al. Accurate design of megadalton-scale two-component icosahedral protein complexes. Science 353, 389–394 (2016).

41 McLellan, J. S., et al. Structural basis of respiratory syncytial virus neutralization by motavizumab. Nat Struct Mol Biol 17, 248–250 (2010).

42 Umemoto, S., et al. Cationic-nanogel nasal vaccine containing the ectodomain of RSV-small hydrophobic protein induces protective immunity in rodents. NPJ Vaccines 8, 106 (2023).

43 Hou, X., Zaks, T., Langer, R. & Dong, Y. Lipid nanoparticles for mRNA delivery. Nat Rev Mater 6, 1078–1094 (2021).

44 Hald Albertsen, C., et al. The role of lipid components in lipid nanoparticles for vaccines and gene therapy. Adv Drug Deliv Rev 188, 114416 (2022).

45 Kwon, D. I., et al. Mucosal unadjuvanted booster vaccines elicit local IgA responses by conversion of pre-existing immunity in mice. Nat Immunol 26, 908–919 (2025).

46 Baldeon Vaca, G., et al. Intranasal mRNA-LNP vaccination protects hamsters from SARS-CoV-2 infection. Sci Adv 9, eadh1655 (2023).

47 Mas, V., Nair, H., Campbell, H., Melero, J. A. & Williams, T. C. Antigenic and sequence variability of the human respiratory syncytial virus F glycoprotein compared to related viruses in a comprehensive dataset. Vaccine 36, 6660–6673 (2018).

48 Zhang, G., Zhao, B. & Liu, J. The Development of Animal Models for Respiratory Syncytial Virus (RSV) Infection and Enhanced RSV Disease. Viruses 16 (2024).

## References and Notes

1 Deng, J. et al. Decoding protective immunity to RSV: A functional epitope landscape prescribes next-generation pediatric vaccine design. bioRxiv (2026).

