## supplemental Fig1-10, Table1-3 for "Rational antigen engineering and mucosal delivery design for next-generation RSV vaccines"

**Materials and Methods**

**Animal ethics and biosafety Statement**

All animal immunization and respiratory syncytial virus (RSV) challenge studies were reviewed and approved by the Experimental Animal Ethics Committee of Sinovac Life Sciences Co., Ltd (202301002, 202406002 and 202506006). Mice and cotton rats were randomly assigned to experimental groups and immunized with the indicated RSV vaccine candidates or control formulations according to the specified schedules. RSV challenge experiments were conducted in a biosafety level 2 (BSL-2) laboratory. All procedures, including immunization, sample collection, viral challenge, and euthanasia, were performed in accordance with institutional guidelines and all relevant ethical regulations for the care and use of laboratory animals.

**Cells and viruses**

HEK293F cells were purchased from Thermo Fisher Scientific and cultured in OPM-293 CD05 Medium (OPM Biosciences) in a 37 °C shaker incubator with 8% CO_2_. HEp-2 (ATCC CCL-23) were cultured in Dulbecco's Modified Eagle Medium (DMEM) supplemented with 10% fetal bovine serum (FBS). RSV strains A2 (VR-1540), Long (VR-26), B9320 (VR-955), and 18537 (VR-1580) were obtained from the American Type Culture Collection (ATCC). RSV was cultured in HEp-2 cells supplemented with 2% FBS and purified via density gradient centrifugation.

**Expression and purification of F mutations and antibodies**

The RSV F, which contains residues 1-513, was cloned into the pCAGGS vector. To produce all proteins, the constructed plasmids were transiently transfected into HEK293F. The cells were maintained at 37 °C in a humidified incubator with 8% CO₂ under agitation at 130 rpm. After 72 hours of post-transfection incubation, the culture was harvested by centrifugation at 1,000 × g for 30 minutes to remove cells. The clarified supernatant was collected, and the target protein was initially purified via affinity chromatography using Strep-Tactin resin. Subsequently, size-exclusion chromatography (SEC) was performed using a Superdex 200 column (GE Healthcare) equilibrated with phosphate-buffered saline (PBS).

The RSV F truncated constructs and Head38-50A constructs were cloned into the pCAGGS vector with a C-terminal His tag. For protein expression, the constructed plasmids were transiently transfected into HEK293F cells. Cells were maintained at 37 °C in a humidified incubator with 8% CO₂ under agitation at 130 rpm. At 72 h post-transfection, the culture supernatant was harvested by centrifugation at 1,000 × g for 30 min to remove cells and debris. The clarified supernatant was collected, and His-tagged target proteins were initially purified by nickel-affinity chromatography using Ni-NTA resin. The eluted proteins were further purified by size-exclusion chromatography using a Superdex 200 column equilibrated with buffer containing 50 mM Tris and 500 mM NaCl.

The 50B protein was cloned into the pET28a(+) vector. For protein expression, the constructed plasmid was transformed into *BL21*. Cells were cultured in LB medium at 37 °C until the optical density at 600 nm reached 0.6–0.8, and protein expression was then induced with IPTG. After 20 hours induction, cells were harvested by centrifugation and resuspended in lysis buffer containing 50 mM Tris, 500 mM NaCl, and 0.75% CHAPS. Cells were lysed by sonication, and the lysate was clarified by centrifugation. The His-tagged 50B protein was purified by nickel-affinity chromatography using Ni-NTA resin. For Head38-50AB nanoparticle assembly, purified Head38-50A and 50B proteins were mixed at a molar ratio of 1:1.2 and incubated overnight at 4 °C to allow particle assembly. The assembled 50AB nanoparticles were purified by size-exclusion chromatography using a Superdex 200 column equilibrated with buffer containing 50 mM Tris and 500 mM NaCl.

All antibodies were synthesized and subcloned into the pcDNA3.1 vector for expression in HEK293F cells. Recombinant plasmids encoding the heavy and light chains of each antibody were transiently co-transfected into HEK293F cells, which were then cultured at 37 °C with 8% CO₂ for 7 days to allow antibody expression. The culture supernatant was harvested and applied to a Protein A CIP column (GenScript) for purification. After washing and elution, the eluate was buffer-exchanged into 1× phosphate-buffered saline (PBS, pH 7.2).

To obtain the Fab fragments of CNR2292 and Motavizumab, the Pierce FAB Preparation Kit (Thermo Scientific) was used. Purified mAbs were incubated with papain and digested at 37 °C for 5 hours. The digested samples were desalted using a desalting column, and the Fab fragments were separated by a Protein A column, which binds the Fc fragment. The purified Fab fragments were then collected for cryo-electron microscopy (cryo-EM) analysis.

**Screening of preF mutations by ELISA**

For supernatant ELISA, MEDI8897 and AM14 were individually coated onto 96-well plates and incubated overnight at 4 °C. The plates were then blocked with PBS containing 1% BSA for 1 h. Serial dilutions of supernatants were added to each well and incubated for 2 h at room temperature. After washing the plates three times with PBST (PBS supplemented with 0.05% Tween-20), mouse anti-strep antibody was added and incubated for 1 h at room temperature. Then goat anti-mouse Fc secondary antibody (1:5000, Abcam) was added and incubated for 1 h at room temperature. The plates were washed five times with PBST, followed by incubation with TMB substrate. The reaction was stopped by adding 50 μL of 2 M H_2_SO_4_, and absorbance at 450 nm was measured using a Thermo Varioskan Flash microplate reader.

For protein ELISA, preF mutants were individually coated onto 96-well plates and incubated overnight at 4 °C. The plates were then blocked with PBS containing 1% BSA for 1 h. Serial dilutions of monoclonal antibodies (mAbs) were added to each well and incubated for 2 h at room temperature. After washing the plates three times with PBST (PBS supplemented with 0.05% Tween-20), HRP-conjugated goat anti-human Fc secondary antibody (1:5000, Abcam) was added and incubated for 1 h at room temperature. The plates were washed five times with PBST, followed by incubation with TMB substrate. The reaction was stopped by adding 50 μL of 2 M H_2_SO_4_, and absorbance at 450 nm was measured using a Thermo Varioskan Flash microplate reader. The half-maximal effective concentration (EC_50_) values of each mAb were determined by nonlinear regression analysis using GraphPad Prism (v10.4.2).

**preF thermal stability assay**

PaSTRy was performed with SYPRO Orange (Invitrogen, Carlsbad, USA) as a fluorescent probe to detect the exposed hydrophobic residues by an MX3005 qPCR instrument (Agilent, Santa Clara, USA). Here, we set up pH = 7.4 25 μL reaction system which contained 10 μg of target protein, 1000x SYPRO Orange, and ramped up the temperature from 25 °C to 99 °C. Fluorescence was recorded in triplicate at an interval of 1 °C.

**Cryo-EM sample preparation and data collection**

The purified RSV F R296 protein was concentrated to 2 mg/mL, followed by the addition of 0.05% N-Dodecyl-β-D-maltoside (DDM). Samples were then dropped onto pre-glow-discharged gold grid (C-flat, 300-mesh, 1.2/1.3), blotted for 6 seconds with no force in 100% relative humidity and immediately plunged into the liquid ethane using Vitrobot (FEI). Cryo-EM data were collected with FEI Titan Krios microscope at 300kV. A total of 6,040 movies were recorded by Falcon4 (total dose of 50.43 e^-^/Å^2^, exposure time of 8.23 s). Automated single particle data acquisition was carried out by EPU, with a calibrated magnification of 75,000 yielding a final pixel size of 1.036 Å.

The truncated RSV F Head38 protein was concentrated to 1.2 mg/mL and incubated with a 1.1-fold molar excess of antibody Fabs (CNR2292 and Motavizumab), followed by the addition of 0.05% N-Dodecyl-β-D-maltoside (DDM). Samples were then dropped onto pre-glow-discharged gold grid (C-flat, 300-mesh, 1.2/1.3), blotted for 6 seconds with no force in 100% relative humidity and immediately plunged into the liquid ethane using Vitrobot (FEI). Cryo-EM data were collected with FEI Titan Krios microscope at 300kV. A total of 4,733 movies were recorded by K3 (total dose of 60 e^-^/Å^2^, exposure time of 3.43 s). Automated single particle data acquisition was carried out by SerialEM, with a calibrated magnification of 22,500 yielding a final pixel size of 1.07 Å.

**Cryo-EM data processing and model building and refinement**

A total of 6,040 movies of R296 were recorded and subjected to patch motion, and defocus value of each motion corrected micrograph was estimated by patch CTF estimation in cryoSPARC ^1^. 2,130,266 particles were picked by template and extracted for 2D classification. 376,946 particles were then processed by heterogeneous refinement. After that, 97,251 particles were selected and processed by nonuniform refinement. The resolution of structure was determined based on the gold-standard Fourier shell correlation (threshold = 0.143) and evaluated by local resolution.

A total of 4,733 movies of Head38 in complex with CNR2292 and Motavizumab were recorded and subjected to patch motion, and defocus value of each motion corrected micrograph was estimated by patch CTF estimation in cryoSPARC ^1^. 2,504,966 particles were picked by template and extracted for 2D classification. 418,871 particles were then processed by heterogeneous refinement. After that, 369,745 particles were selected and processed by nonuniform refinement. The resolution of structure was determined based on the gold-standard Fourier shell correlation (threshold = 0.143) and evaluated by local resolution.

The previously determined RSV F structure (4MMT) was used to generate the RSV pre-F trimer model and Head38 model. PDBs were fitted into the cryo-EM densities by Chimera. Manual adjustment and correction according to protein sequences and densities use Coot and real space refinement use Phenix.

**Animal experiments**

All animals were immunized according to the experimental schedule described in this study. After optimization of the immunization regimen, a 0/21-day prime–boost schedule was selected for all immunization experiments. All protein vaccines were formulated with Al(OH)₃ adjuvant.

BALB/c mice were bled on day 35, and spleens were collected after euthanasia. SD rats were bled on days 35, 49, 63, 91, and 119. Cotton rats were bled on days 35 and 49, and were intranasally challenged with RSV A2 on day 49. Each cotton rat received 1 × 10⁶ PFU virus in a total volume of 100 μL. Animals were euthanized on day 54, and lung tissues and nasal turbinates were collected.

Blood samples were incubated at 37 °C for 60 min, followed by centrifugation at 3000 g for 15 min at 4 °C to separate serum. The collected sera were used for RSV neutralization assays and preF-binding antibody detection.

For lung tissues, the left lung was used for H&E staining and pathological evaluation, while the right lung was used for viral load quantification and cytokine expression analysis.

**Microneutralization Assay for Serum Neutralizing Antibodies**

Except cotton rats immunized with the mRNA vaccine and SD rats for cross-neutralization assays, a luciferase-based neutralization assay was used to determine serum neutralizing antibody titers. The virus used in the assay was RSV-Long-Luc. Serum samples were serially diluted in 96-well plates and mixed with RSV-Long-Luc virus. The mixtures were incubated at 37 °C in a 5% CO₂ incubator for 1 h to allow virus neutralization. After incubation, 293T cells were added to each well and the plates were further incubated at 37 °C with 5% CO₂ for 48 h. After 48 h, luciferase substrate was added to each well and luminescence (RLU) was measured using a microplate reader. The neutralizing antibody titer (NT_50_) was defined as the reciprocal of the serum dilution that inhibited 50% of RSV infection.

For cotton rat sera and for cross-neutralization assays of SD rat sera against multiple RSV strains, serum samples were serially diluted in 96-well plates and mixed with RSV virus, followed by incubation at 37 °C with 5% CO₂ for 2 h. HEp-2 cells were then added and plates were incubated at 37 °C with 5% CO₂ for 5 days. After incubation, the supernatant was removed and cells were fixed with 80% ice-cold acetone at 4 °C for 15 min. After fixation, 75 μL of 5% BSA was added to each well for blocking at room temperature for 1 h. Plates were washed three times with TBST, followed by incubation with anti-RSV rabbit polyclonal antibody at 37 °C for 1 h. After washing, HRP-labeled goat anti-rabbit IgG was added and incubated at 37 °C for 1 h. After washing, 100 μL TMB substrate was added to each well and incubated at room temperature for 10 min. The reaction was stopped by adding 1% HCl, and absorbance was measured at 450 nm. Neutralizing antibody titers (NT_50_) were defined as the reciprocal of the serum dilution that inhibited 50% RSV infection.

**ELISA for Serum Binding IgG Antibodies**

The ELISA method was used to determine preF-binding antibody titers in cotton rat sera. Briefly, 96-well plates were coated with 50 μL RSV preF antigen (2.0 μg/mL) and incubated at 2–8 °C overnight. The coated plates were washed once and then blocked with 100 μL blocking buffer at room temperature for 1.5 h. After washing, serum samples were 5-fold serially diluted (8 dilution points) and 50 μL diluted serum was added to each well, followed by incubation at room temperature for 2 h. After washing three times, Chicken anti-cotton rat IgG (H&L) secondary antibody conjugated with HRP diluted 1:5000 was added (50 μL per well) and incubated at room temperature for 45 min in the dark. After washing, TMB substrate (mixed 1:1) was added (50 μL per well) and incubated in the dark for 10 min. The reaction was stopped by adding 50 μL stop solution, and absorbance was measured at 450 nm using a microplate reader.

**ELISPOT assay**

Cellular immune responses in BALB/c mice were evaluated using an ELISPOT assay. Mice were euthanized by cervical dislocation and spleens were collected. Splenocytes were obtained by mechanical disruption, followed by red blood cell lysis and centrifugation at 500 g for 5 min. Cells were resuspended in RPMI 1640 complete medium containing 10% FBS, counted, and adjusted to the required concentration. Cells were cultured at 37 °C with 5% CO₂. In the ELISPOT plates, 2.5 or 2 × 10⁵ splenocytes per well were added. The RSV F peptide pool was used at a final concentration of 2 μg/mL, and ConA was used as a positive control at 1 μg/mL. The assay was performed according to the standard ELISPOT protocol.

**Plaque Assay for Viral Loads in Lung and Nasal Turbinates**

Lung viral loads were determined by plaque assay. Briefly, the right lungs were collected and homogenized, and the homogenates were titrated on HEp-2 cells in 24-well plates. After infection for 2 h at 37 °C, the inoculum was removed and cells were overlaid with DMEM containing 1.2% methylcellulose and 2% FBS. Plates were incubated at 37 °C with 5% CO₂ for 7 days. Cells were then fixed and stained with 0.2% crystal violet solution, and plaques were counted under a microscope by an investigator blinded to the experimental conditions. Viral titers were calculated and expressed as PFU per gram of lung tissue (PFU/g). The limit of detection was 25 PFU/g of tissue.

**qPCR Analysis of Pulmonary Cytokine Expression**

Total RNA was extracted from lung tissues using a Zymo Research RNA extraction kit according to the manufacturer’s protocol following tissue homogenization. cDNA was synthesized using a Takara reverse transcription kit according to the manufacturer’s instructions.

cDNA generated from lung tissue RNA was used as the template for cytokine expression analysis. Quantitative PCR was performed using ChamQ Blue Universal SYBR qPCR Master Mix (Vazyme) with gene-specific primers.

Relative expression levels of cytokine genes were calculated using the 2^-ΔΔCt^ method, with β-actin as the internal reference gene. Gene expression levels were normalized to the control group and presented as relative fold changes.

**Histopathological Evaluation of Lung Inflammation by H&E Staining**

At day 54, lung tissues were collected and fixed in 4% paraformaldehyde for more than 24 h. The fixed tissues were dehydrated through a graded ethanol series, cleared in xylene, embedded in paraffin, and sectioned at a thickness of 4 μm. Tissue sections were deparaffinized, rehydrated, and stained with hematoxylin and eosin (H&E) following standard procedures, followed by dehydration and mounting. Histopathological changes in lung tissues were evaluated using a semi-quantitative scoring system. Four parameters were assessed, including inflammatory cell infiltration in the alveolar region, perivascular region, peribronchiolar region, and interstitial tissue ^2^. Each parameter was scored on a scale of 0–4 based on the severity of inflammation: 0, no visible lesion; 1, mild; 2, moderate; 3, severe; and 4, very severe.

**mRNA pharmacokinetic and tissue distribution analysis**

To evaluate the pharmacokinetic and tissue distribution profiles of the lyophilized RSV mRNA vaccine, EDTA-anticoagulated whole blood and major tissues, including brain, heart, liver, spleen, lung, kidney, and muscle, were collected from SD rats at predefined time points after administration. Samples were stored at −60 °C or below until total RNA extraction. RSV mRNA levels were quantified using a validated RT-qPCR assay targeting a specific sequence within the RSV mRNA vaccine. Unencapsulated RSV mRNA vaccine was used as the reference standard to generate calibration curves, and the mRNA amount in each reaction was back-calculated from the quantification cycle (Cq) values. The assay had a quantitative range of 5.00 × 10⁻⁴ to 1.00 × 10² pg/reaction, with calibration curve R² values of 0.998–1.000 and amplification efficiencies of 92.5%–96.0%. Method validation demonstrated acceptable precision, accuracy, selectivity, matrix effect, dilution linearity, recovery, and stability. The final results were expressed as the amount of RSV mRNA per microgram of total RNA and used for subsequent pharmacokinetic and tissue distribution analyses.

**Lipid pharmacokinetic and tissue distribution analysis**

To evaluate the pharmacokinetic and tissue distribution profiles of the lipid components of the RSV mRNA vaccine, plasma and major tissue samples were collected from SD rats at predefined time points after administration. Tissue samples, including brain, gonads, muscle, thymus, heart, lung, kidney, liver, spleen and lymph nodes, were homogenized with dilution buffer and stored at −20 °C or below until analysis. The concentrations of SM-102 and mPEG-DMG-2K in rat plasma and tissue homogenates were quantified using a validated LC-MS/MS method, with ALC-0315 used as the internal standard. Calibration standards and quality control samples were prepared in blank rat plasma or blank tissue homogenate matrices and processed according to the validated sample preparation procedure. For tissue samples, the validated quantitative ranges were 50–25,000 ng/g for SM-102 and 250–125,000 ng/g for mPEG-DMG-2K. For plasma samples, the quantitative ranges were 1–500 ng/mL for SM-102 and 5–2,500 ng/mL for mPEG-DMG-2K. Method validation demonstrated acceptable linearity, precision, accuracy, system carryover, dilution reliability, extraction recovery, injection reproducibility and sample stability. LC-MS/MS data were acquired using Analyst software and processed with Watson LIMS. Concentrations of SM-102 and mPEG-DMG-2K were calculated from calibration curves based on the peak-area ratios of analyte to internal standard. Final results were expressed as ng/mL for plasma and ng/g for tissues and used for subsequent lipid pharmacokinetic and tissue distribution analyses.

**Preparation and characterization of LNP**

LNPs were prepared with a microfluidic device, ionizable lipids, cholesterol, DSPC, cationic lipid and DMG-PEG2000 (40:37.5:20:2:1.5 and 50:38.5:10:0:1.5) were dissolved in ethanol to form representative LNP. mRNA was diluted in sodium citrate buffer solution (pH = 4.0, 10 mM). The two substances were mixed in a microfluidics chip with a volume ratio of 1/3 (flow rate was 12 mL/min), and the N/P was 4.5. The LNPs were dialyzed in PBS buffer (pH = 7.8, 0.01 M) with dialysis tubing (MWCO: 10 kDa) for 12 h to obtain the final product.

**Size, PDI, and Zeta Potential**

The particle size, PDI, and zeta potential were determined by dynamic light scattering (DLS). The prepared LNP solution was diluted 100 times with deionized water, then tested with the DTS1070 sample cell at room temperature.

**Encapsulation Efficiency**

The encapsulation efficiency of mRNA in LNP was determined by using Thermo Fisher’s Quant-iT RiboGreen RNA Quantification kit. The fluorescence intensity was measured under the condition of excitation light with a wavelength of 480 nm and emission light with a wavelength of 530 nm by a microplate reader. The fluorescence

intensity of the sample was converted into the corresponding mRNA concentration in the standard curve. Encapsulation efficiency formula = (total mRNA concentration- free mRNA concentration) / total mRNA concentration of demulsified sample ×100%.

***In vivo* bioluminescence**

Female BALB/c mice (6–8 weeks) were intranasally administered LNP containing 2 μg of Luciferase mRNA (APExBIO). 6 h after administration, mice were intraperitoneally injected with 200 μL of 150 mg/kg D-luciferin. 5 mins later, using IVIS Spectrum 2 (Revvity, Shanghai) measure the bioluminescence intensity at a wavelength of 540 to 600 nm.

Female BALB/c mice (6–8 weeks) were intranasally administered with LNP containing 2 μg of Luciferase mRNA (APExBIO). 0, 3, 6, 9, 12, 15 hours post-administration, mice were intraperitoneally injected with 200 μL of 150 mg/kg D-luciferin. 5 mins later, using IVIS Spectrum 2 (Revvity, Shanghai) measure the fluorescence intensity at a wavelength of 540 to 600 nm

**Cryogenic transmission electron microscopy (cryoTEM)**

An aliquot of 4 μL lipid nanoparticle solution was applied onto the glow-discharged Quantifoil Cu R1.2/1.3 400 mesh grid or Xinxing Bairui T11032 ultra-thin carbon supported grids, waited for 60s, blotted with filter paper for 3 s, and plunge-frozen into liquid ethane using a Thermo Fisher Vitrobot Mark IV. Cryo-EM data acquisition was performed on a 300 kV Thermo Fisher Krios G4 electron microscope equipped with a Falcon 4 direct detection camera and a Selectris energy filter (GIF: a slit width of 10 eV), at counting mode (bin2). The micrographs were collected at a calibrated magnification of 37,000×, corresponding to a pixel size of 0.48 nm; 59,000×, coresponding to a pixel size of 0.31 nm and 75,000×, coresponding to a pixel size of 0.24 nm, respectively. In summary, a total of 15 micrographs were collected on single frame micrograph. Data acquisition parameters can be found in the table above.

**Scanning electron microscopy (SEM)**

Nasal mucosal tissues were taken around 80 nm using an UC7 (Leica, Wetzlar, Germany) picked up on formvar/Carbon coated 100 mesh Cu grids, stained for 40s in 3.5% Uranyl Acetate in 50% Acetone followed by staining in Sato’s Lead Citrate for 2min.Observed in the JEOL JEM-1400 120kV. Images were taken using a Gatan OneView 4k X 4k digital camera. For scanning electron microscopy analysis. Samples were then dehydrated in a series of ethanol. Then dried in a Critical Point Dryer, Sputter coated (Leica EM ACE600) with gold. Observed in Zeiss Sigma FESEM. Images were taken using Smart SEM VO5.03.

**Mouse immunization for LNP**

For immunization, mice were intranasally administered 20 μL of LNP containing 10 μg RSV antigen mRNA with CpG co-encapsulated in the LNP per mouse. At various point post-immunization, samples from blood, nasal lavage fluid (NLF), bronchoalveolar lavage fluid (BALF) and spleens were collected from each group, spleens were processed into single-cell suspensions for ELISpot analysis, and the levels of antigen-specific antibodies and cellular immune responses were assessed. Female mice were used in this initial immunogenicity study due to their generally enhanced immune responsiveness to vaccines and to maintain consistency with translational development pipelines.

**Statistical analysis**

Statistical analyses were performed using GraphPad Prism 10.4.2. For comparisons among multiple groups, one-way ANOVA was used. Differences were considered statistically significant when p < 0.05. Significance levels are indicated as follows: ns, p > 0.05; *p < 0.05; **p < 0.01; ***p < 0.001; ****p < 0.0001.

**Data, code, and materials availability:**

All the cryo-EM density maps and corresponding atomic models have been deposited in the Electron Microscopy Data Bank (EMDB, http://www.ebi.ac.uk/pdbe/emdb/) and Protein Bank (PDB, https://www.ebi.ac.uk/pdbe/). The accession codes are available for R296 (EMDB: EMD-81934, PDB: 43KA), Head38 in complex with CNR2292 and motavizumab (EMDB: EMD-81935, PDB: 43KB). The code and data used in this study is available from the corresponding author upon reasonable request.

**References and Notes**

1	Punjani, A., Rubinstein, J. L., Fleet, D. J. & Brubaker, M. A. cryoSPARC: algorithms for rapid unsupervised cryo-EM structure determination. *Nat Methods* **14**, 290-296 (2017).

2	Blanco, J. C., Boukhvalova, M. S., Pletneva, L. M., Shirey, K. A. & Vogel, S. N. A recombinant anchorless respiratory syncytial virus (RSV) fusion (F) protein/monophosphoryl lipid A (MPL) vaccine protects against RSV-induced replication and lung pathology. *Vaccine* **32**, 1495-1500 (2014).

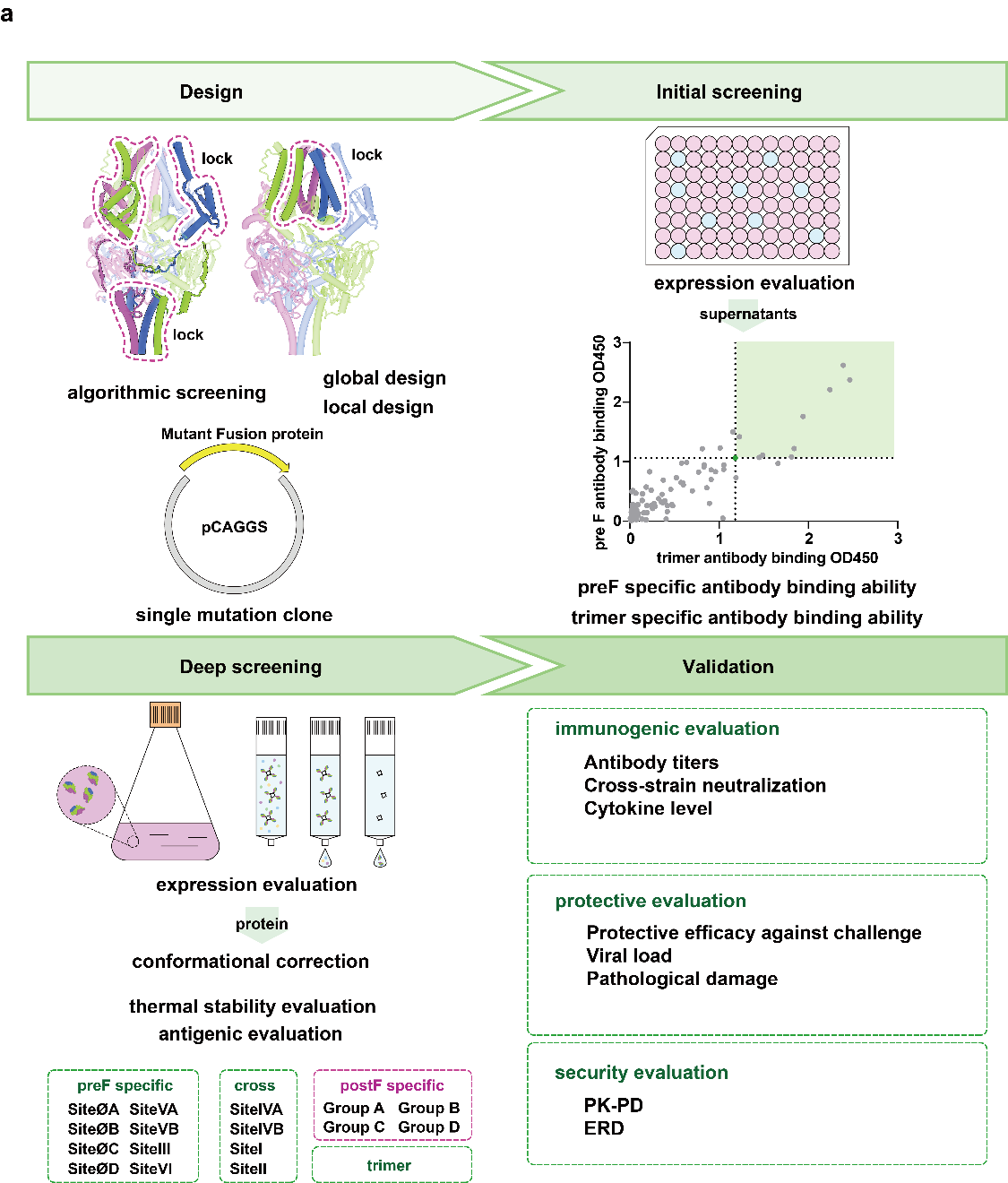

**Extended Data Fig. 1. Flowchart of the four-step screening workflow for RSV F immunogen design. a,** The schematic summarizes the design and selection pipeline, including computational structure-guided engineering, expression and antibody-binding screening, biochemical and antigenic characterization, and animal immunogenicity testing. Major evaluation metrics are indicated for each step.

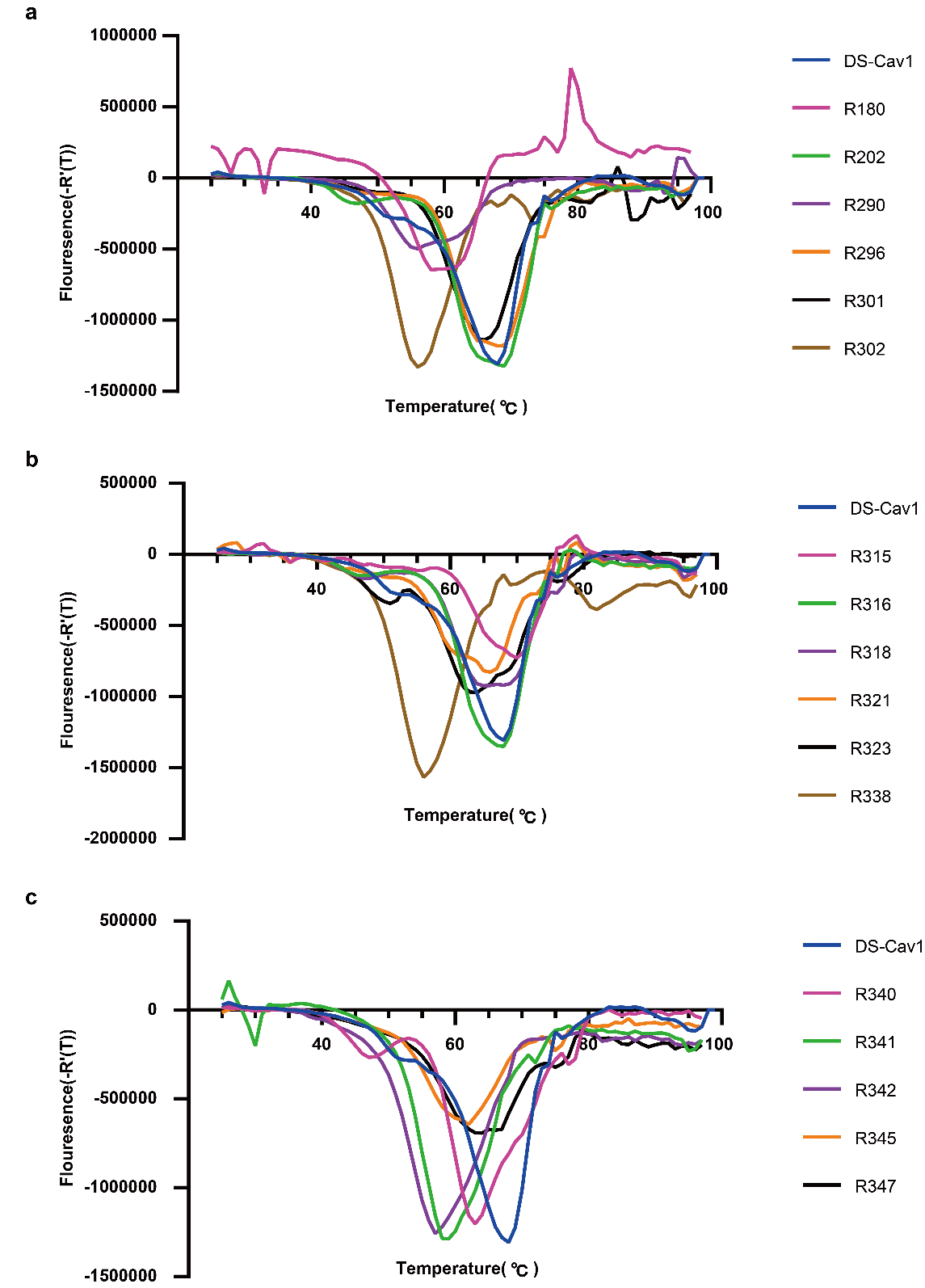

**Extended Data Fig. 2. Raw Thermofluor data used to determine the melting temperatures of RSV F immunogen candidates.**

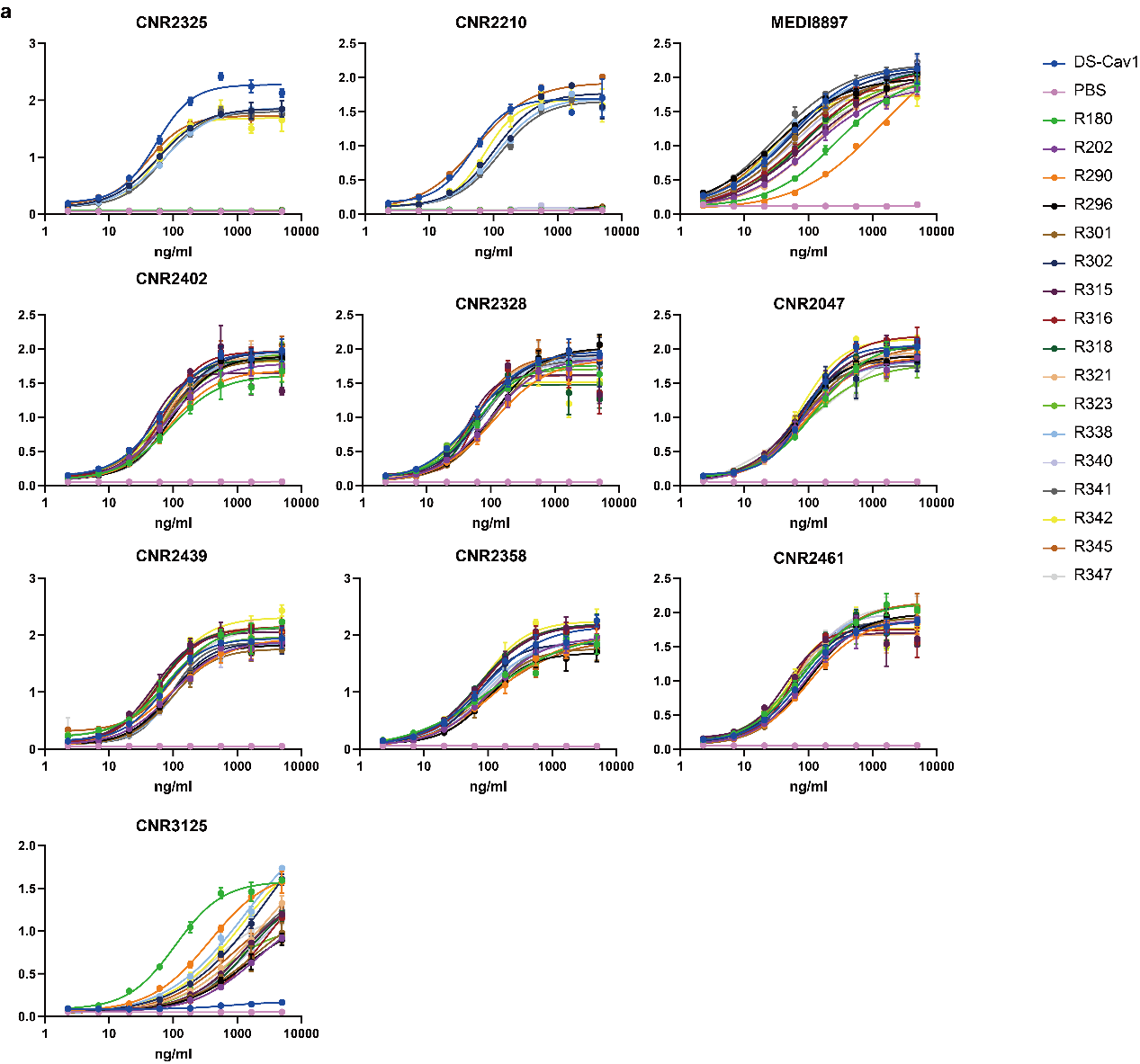

**Extended Data Fig. 3. ELISA binding curves of antibodies against different RSV F antigens.**

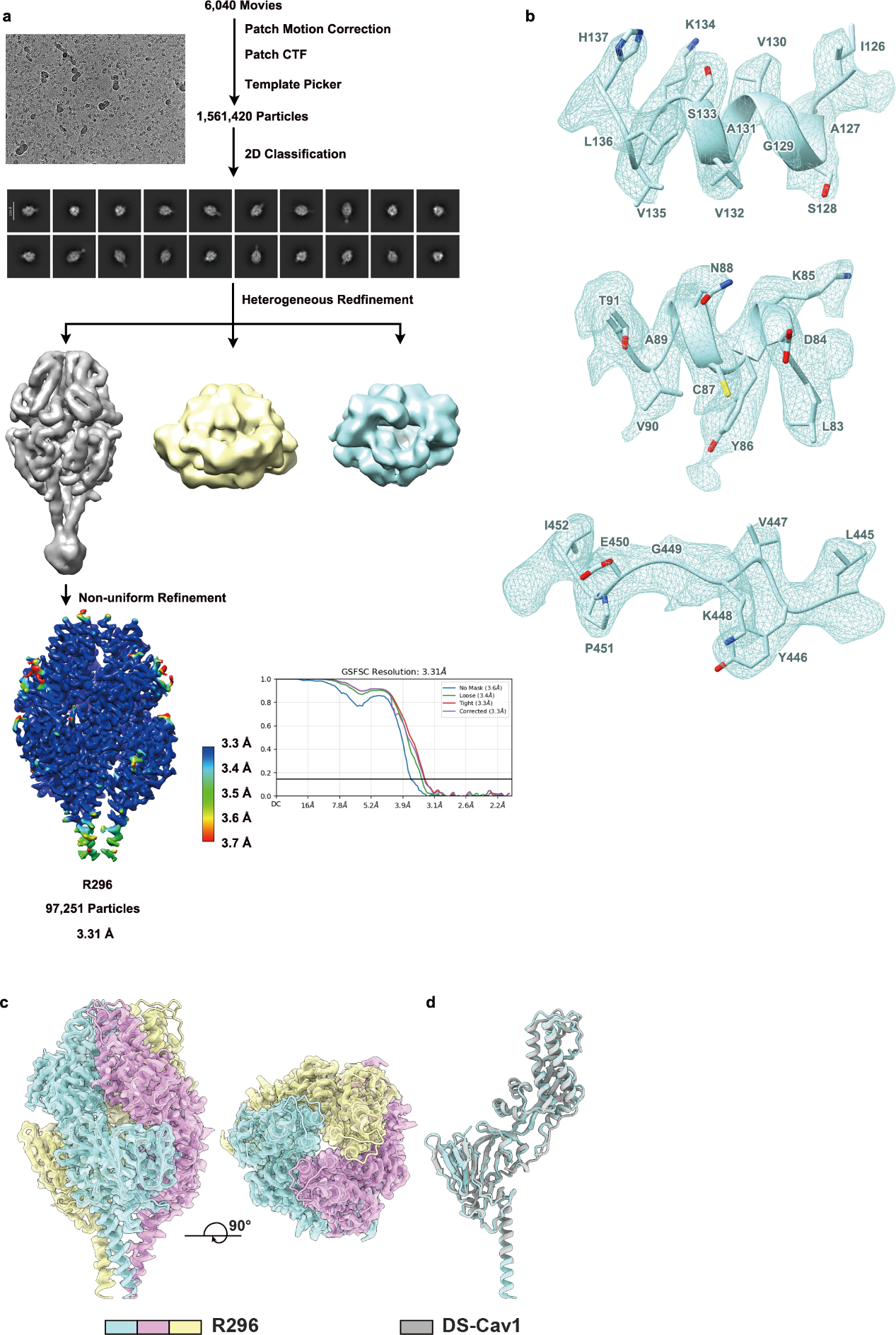

**Extended Data Fig. 4. Cryo-EM structure determination and validation of the R296 immunogen. a,** Cryo-EM data processing workflow and gold-standard Fourier shell correlation (FSC) curves for the final reconstruction. **b,** Local density maps showing representative regions of the cryo-EM reconstruction. **c,** Representative cryo-EM density fitted with the atomic model. **d,** Cartoon comparison of the R296 cryo-EM structure with DS-Cav1.

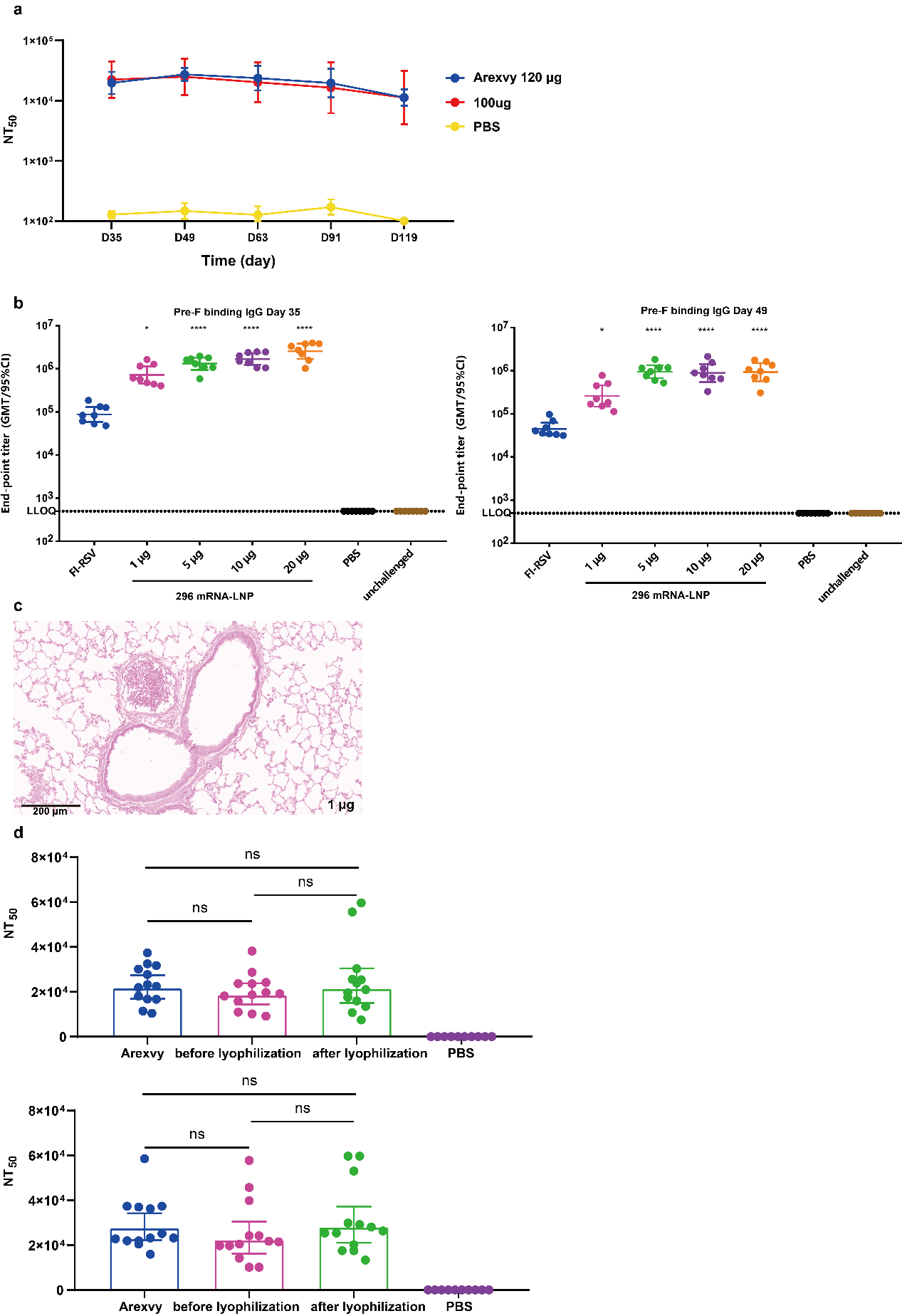

**Extended Data Fig. 5. Additional immunogenicity and histopathological evaluation of R296 mRNA vaccination. a,** Longitudinal serum antibody titers in SD rats measured on days 35, 49, 63, 91, and 119 after R296 mRNA immunization (n = 10). **b,** Serum preF-specific IgG endpoint titers in cotton rats measured on days 35 and 49 after immunization (n = 8). **c,** Representative H&E-stained lung sections from cotton rats immunized with 1 μg R296 mRNA after RSV A2 challenge, showing pulmonary histopathological changes (magnification, ×144; scale bars, 200 μm). **d,** Serum NT_50_ titers induced by R296 mRNA before and after lyophilization in SD rats were compared with Arexvy and PBS controls. Low-dose(50 μg) and high-dose(100 μg) groups are shown in the up and down panels (n = 13), respectively.

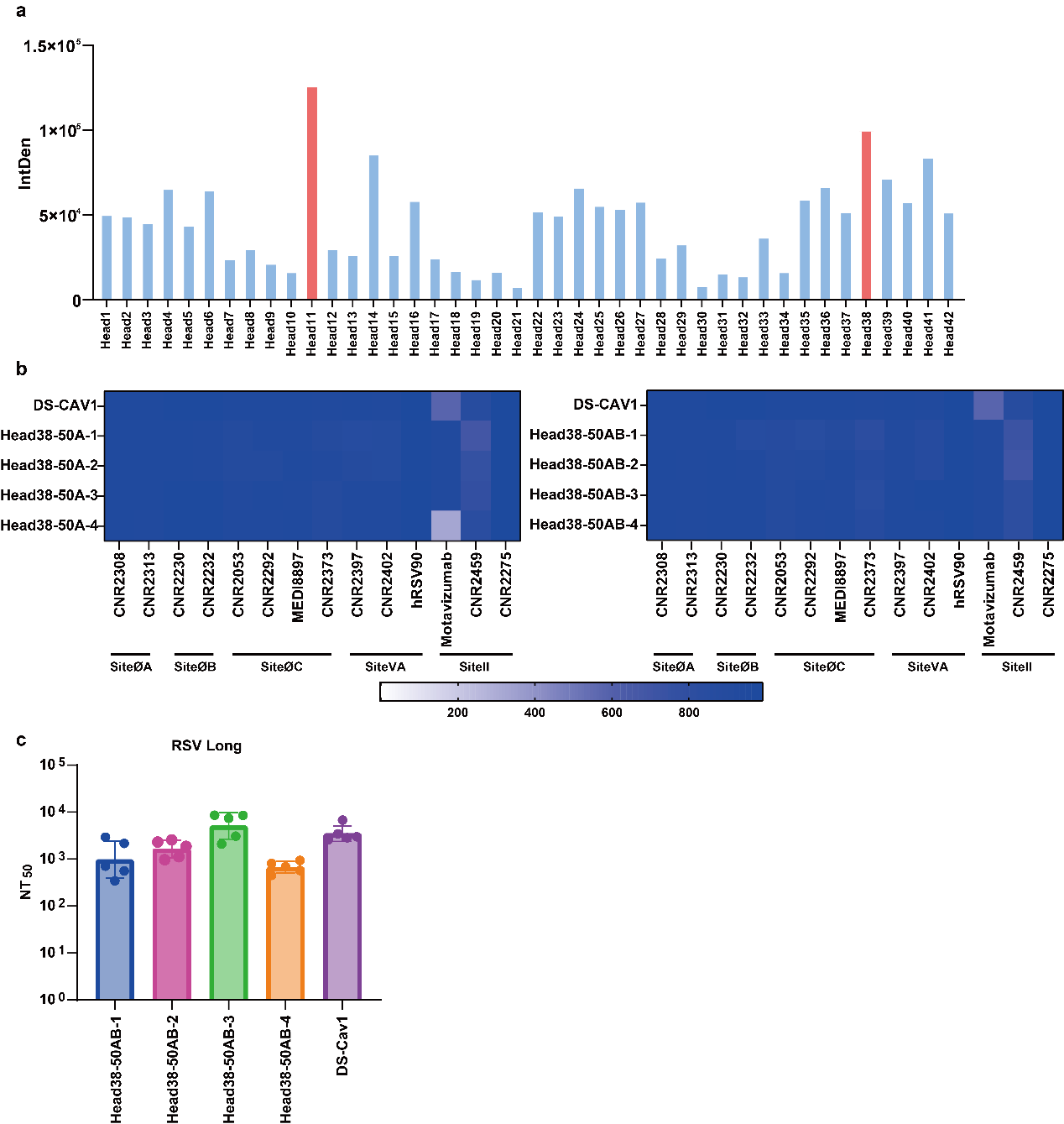

**Extended Data Fig. 6. Screening and characterization of truncated RSV F immunogens displayed on I53-50AB nanoparticles. a,** Relative expression levels of different truncated RSV F constructs quantified by western blot densitometry. **b,** ELISA analysis of I53-50AB nanoparticle-displayed antigens before and after assembly. **c,** Serum NT_50_ titers in mice immunized with nanoparticle immunogens containing different linker lengths (n = 5).

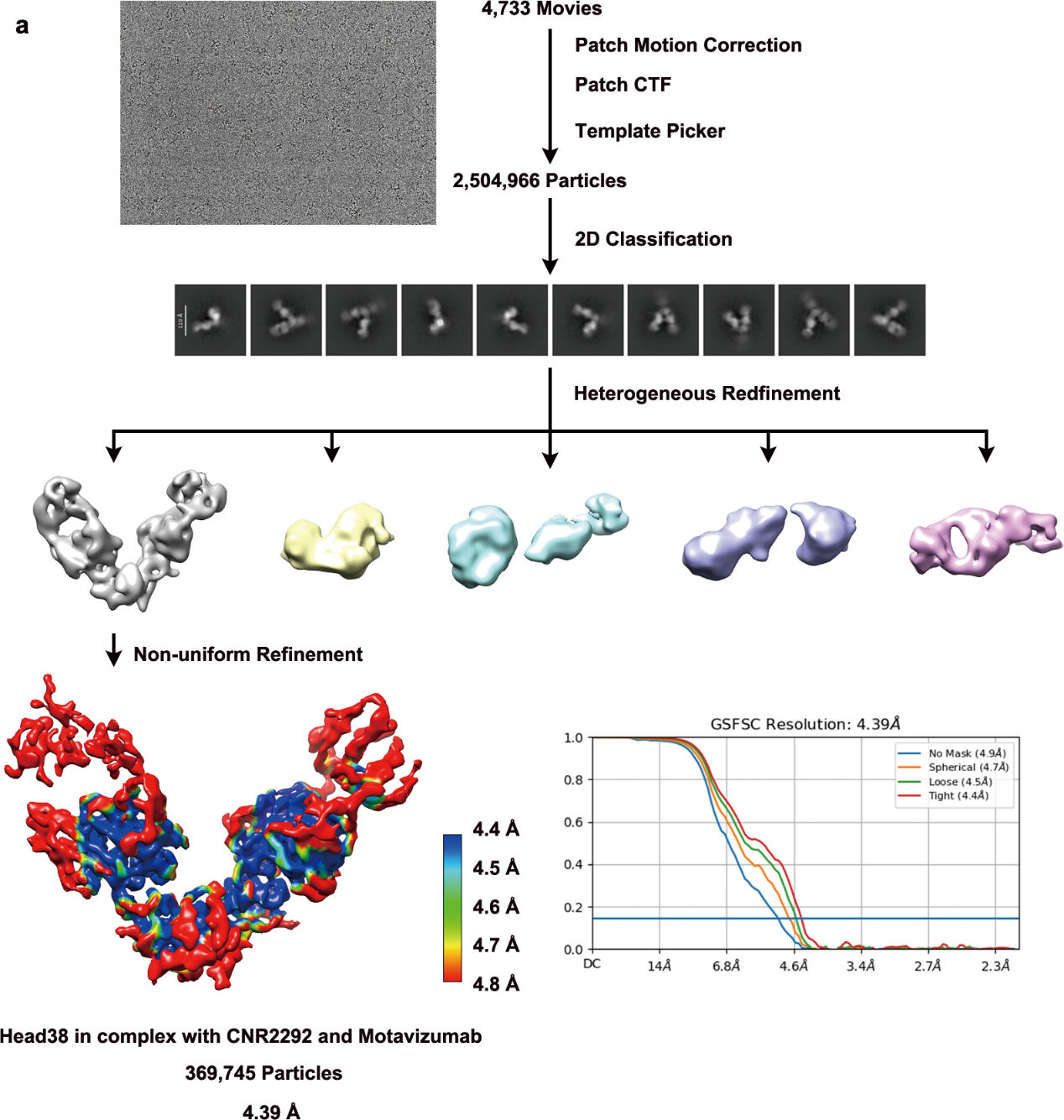

**Extended Data Fig. 7. Cryo-EM structure determination and validation of the Head38 in complex with CNR2292 and Motavizumab. a,** Cryo-EM data processing workflow and gold-standard Fourier shell correlation (FSC) curves for the final reconstruction.

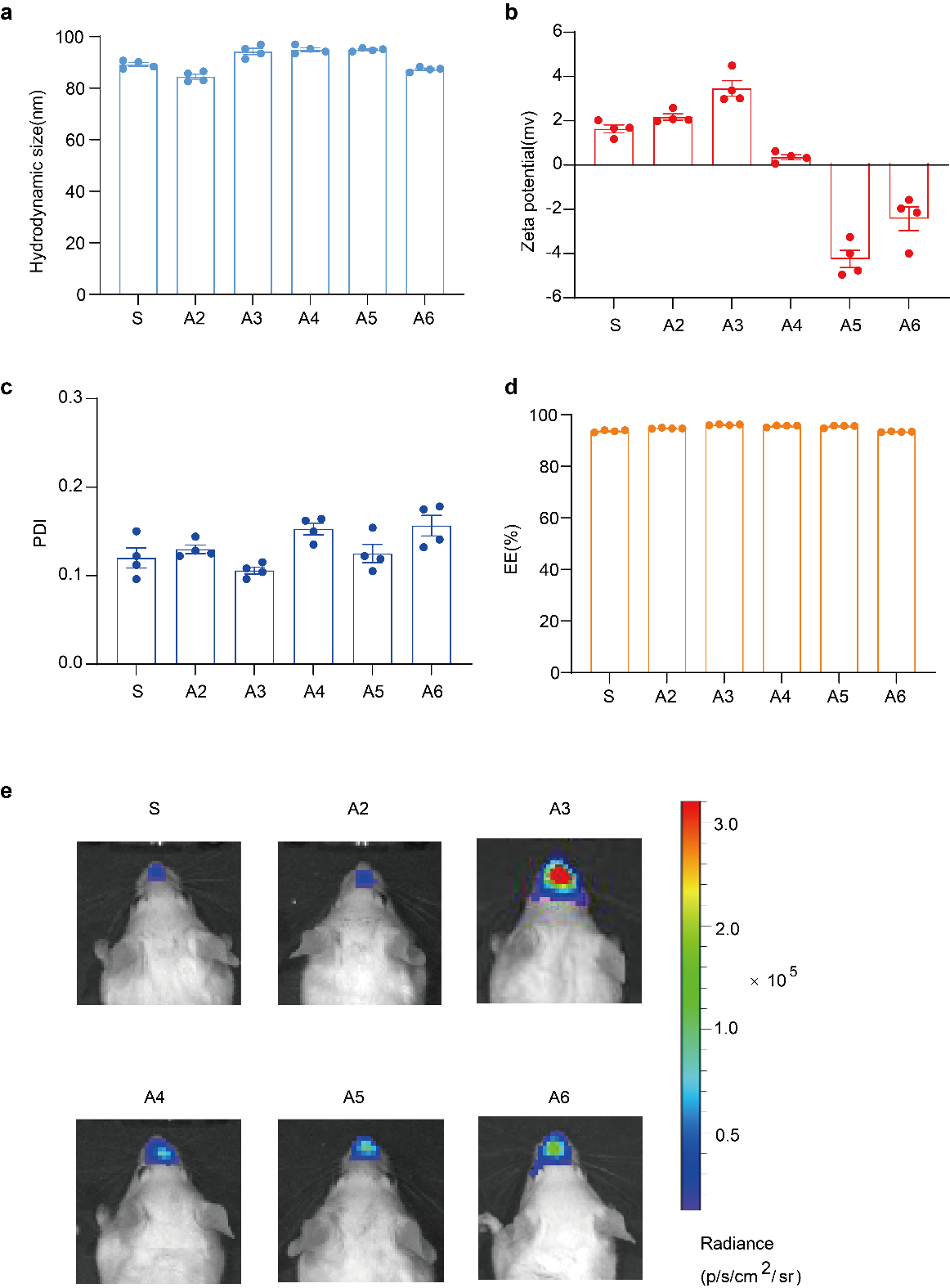

**Extended Data Fig. 8. Physicochemical characterization of six LNP formulations related to Fig. 5a-c. a,b,** Hydrodynamic diameter and zeta potential of the six LNP formulation groups (A1–A6). **c,d,** Polydispersity index (PDI) and encapsulation efficiency (EE%) of the six LNP formulation groups. All formulations showed comparable nanoscale particle sizes, near-neutral surface charge, low PDI values, and high mRNA encapsulation efficiency. Data are shown as mean ± s.e.m. or as individual values (n = 4). **e,** *In vivo* bioluminescence imaging of the nasal cavity in BALB/c mice 6 h after intranasal administration of LNPs containing 2 μg luciferase mRNA. Data are shown for individual animals or as mean ± SEM (n = 4).

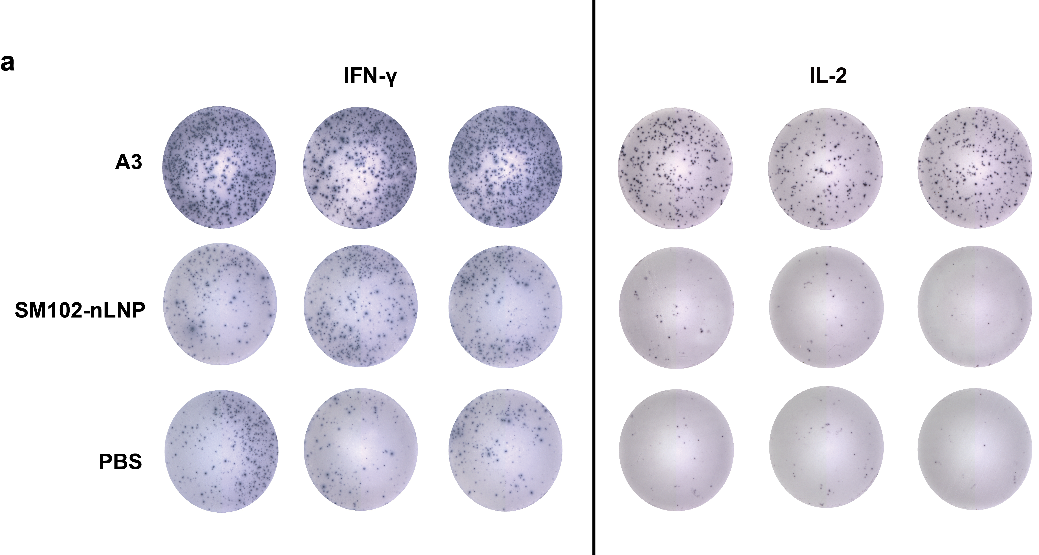

**Extended Data Fig. 9. Cellular immune responses after RSV immunization. a,** IFN-γ and IL-2 ELISpot responses in splenocytes collected on day 42 after primary immunization and restimulated with RSV overlapping peptide pools.

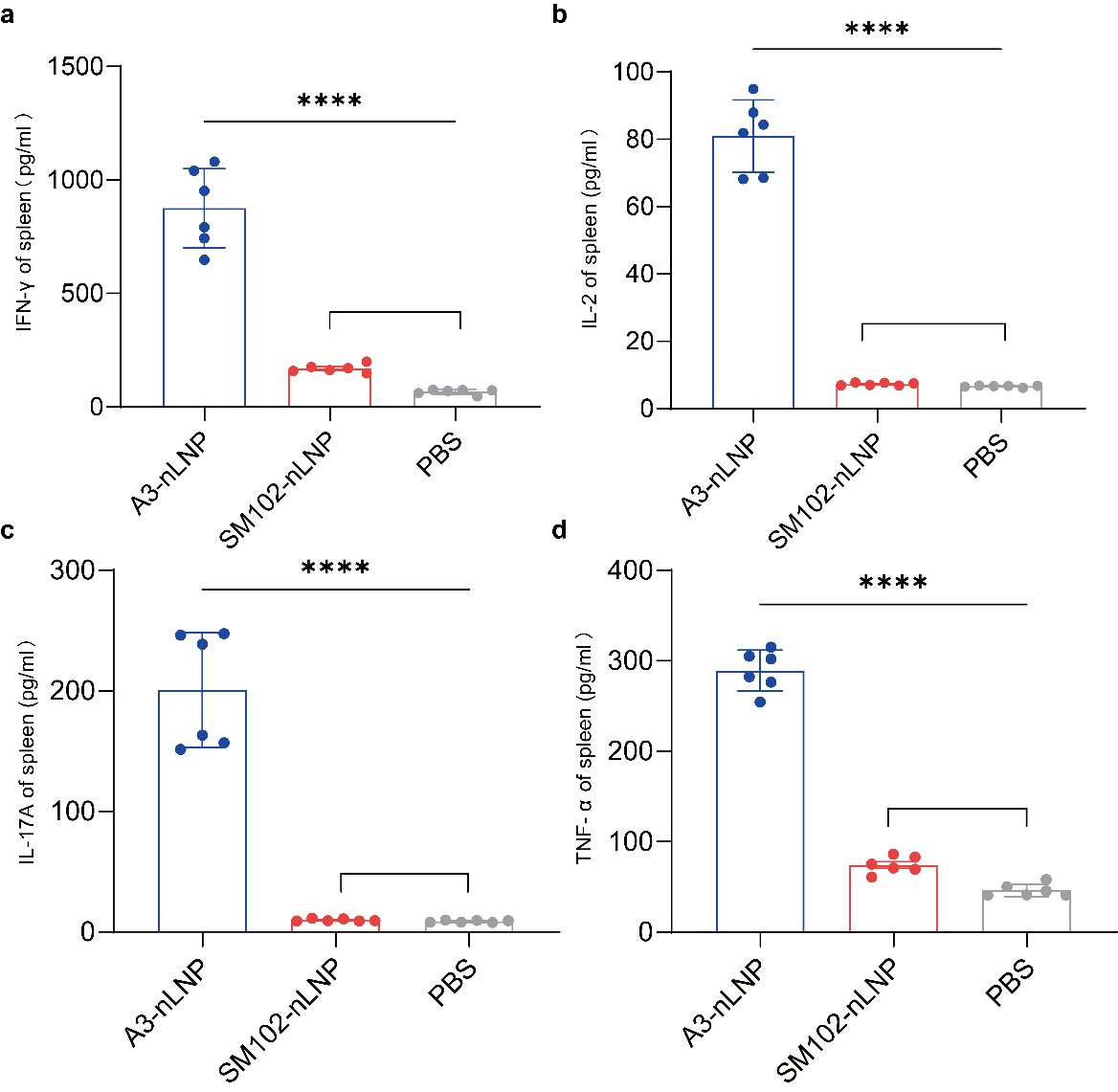

**Extended Data Fig. 10. Cytokine production by splenocytes after RSV antigen restimulation, related to Fig. 5l-m.** Splenocytes were collected from immunized mice on day 42 after primary immunization and restimulated with RSV overlapping peptide pools for 24 h. Cytokine levels in the culture supernatants were measured by ELISA, including IFN-γ **(a)**, IL-2 **(b)**, IL-17A **(c)**, and TNF-α **(d)**. Data are shown as individual animals with group summaries where applicable (n = 6 per group).

**Extended Data Table 1. Design and mutation information of preF constructs.**

| **index** | **mutation site** | **Route 1** | **Route 2** | **Route 3** | **Route 4** | **Route use** |
| --- | --- | --- | --- | --- | --- | --- |
| R024 | DS-CAV1 |  |  |  |  |  |
| R026 | WT |  |  |  |  |  |
| R027 | P484C, I499C | P484C, I499C |  |  |  | 1 |
| R028 | V482C, I499C | V482C, I499C |  |  |  | 1 |
| R029 | V482C, S502C | V482C, S502C |  |  |  | 1 |
| R030 | M396C, F488C | M396C, F488C |  |  |  | 1 |
| R031 | S398C, S485C | S398C, S485C |  |  |  | 1 |
| R032 | S398C, D486C | S398C, D486C |  |  |  | 1 |
| R033 | I148C, Y286C |  | I148C, Y286C |  |  | 2 |
| R034 | H159C, S290C |  | H159C, S290C |  |  | 2 |
| R035 | H159C, I291C |  | H159C, I291C |  |  | 2 |
| R036 | L158C, S290C |  | L158C, S290C |  |  | 2 |
| R037 | I57C, S190C, S180C, S186C |  | I57C, S190C, S180C, S186C |  |  | 2 |
| R038 | I57C, S190C, V178C, V187C |  | I57C, S190C, V178C, V187C |  |  | 2 |
| R039 | I57C, S190C, V178C, L188C |  | I57C, S190C, V178C, L188C |  |  | 2 |
| R040 | I59C, L193C, S180C, S186C |  | I59C, L193C, S180C, S186C |  |  | 2 |
| R041 | I59C, S190C, V178C, V187C |  | I59C, S190C, V178C, V187C |  |  | 2 |
| R042 | I59C, S190C, V178C, L188C |  | I59C, S190C, V178C, L188C |  |  | 2 |
| R043 | Q494C, K399C | Q494C, K399C |  |  |  | 1 |
| R044 | D489C, S398C | D489C, S398C |  |  |  | 1 |
| R045 | D489C, D486C | D489C, D486C |  |  |  | 1 |
| R046 | D489C, S485C | D489C, S485C |  |  |  | 1 |
| R047 | N460C, S146C |  | N460C, S146C |  |  | 2 |
| R048 | A149C, Y458C |  | A149C, Y458C |  |  | 2 |
| R050 | K80A |  |  | K80A |  | 3 |
| R051 | K80I |  |  | K80I |  | 3 |
| R052 | K80L |  |  | K80L |  | 3 |
| R053 | K80V |  |  | K80V |  | 3 |
| R054 | K80F |  |  | K80F |  | 3 |
| R055 | K80W |  |  | K80W |  | 3 |
| R056 | K80Y |  |  | K80Y |  | 3 |
| R057 | K80M |  |  | K80M |  | 3 |
| R058 | K80S |  |  | K80S |  | 3 |
| R059 | K80P |  |  | K80P |  | 3 |
| R060 | N67A, K80A |  |  | N67A, K80A |  | 3 |
| R061 | N67A, K80I |  |  | N67A, K80I |  | 3 |
| R062 | N67A, K80L |  |  | N67A, K80L |  | 3 |
| R063 | N67A, K80V |  |  | N67A, K80V |  | 3 |
| R064 | N67A, K80F |  |  | N67A, K80F |  | 3 |
| R065 | N67A, K80W |  |  | N67A, K80W |  | 3 |
| R066 | N67A, K80Y |  |  | N67A, K80Y |  | 3 |
| R067 | N67A, K80M |  |  | N67A, K80M |  | 3 |
| R068 | N67A, K80S |  |  | N67A, K80S |  | 3 |
| R069 | N67A, K80P |  |  | N67A, K80P |  | 3 |
| R070 | N67A, K87A |  |  | N67A, K87A |  | 3 |
| R071 | N67A, K87I |  |  | N67A, K87I |  | 3 |
| R072 | N67A, K87L |  |  | N67A, K87L |  | 3 |
| R073 | N67A, K87V |  |  | N67A, K87V |  | 3 |
| R074 | N67A, K87F |  |  | N67A, K87F |  | 3 |
| R075 | N67A, K87W |  |  | N67A, K87W |  | 3 |
| R076 | N67A, K87Y |  |  | N67A, K87Y |  | 3 |
| R077 | N67A, K87M |  |  | N67A, K87M |  | 3 |
| R078 | N67A, K87S |  |  | N67A, K87S |  | 3 |
| R079 | N67A, K87P |  |  | N67A, K87P |  | 3 |
| R080 | N67A, K80A, K87A |  |  | N67A, K80A, K87A |  | 3 |
| R081 | N67A, K80A, K87I |  |  | N67A, K80A, K87I |  | 3 |
| R082 | N67A, K80A, K87L |  |  | N67A, K80A, K87L |  | 3 |
| R083 | N67A, K80A, K87V |  |  | N67A, K80A, K87V |  | 3 |
| R084 | N67A, K80A, K87F |  |  | N67A, K80A, K87F |  | 3 |
| R085 | N67A, K80A, K87W |  |  | N67A, K80A, K87W |  | 3 |
| R086 | N67A, K80A, K87Y |  |  | N67A, K80A, K87Y |  | 3 |
| R087 | N67A, K80A, K87M |  |  | N67A, K80A, K87M |  | 3 |
| R088 | N67A, K80A, K87S |  |  | N67A, K80A, K87S |  | 3 |
| R089 | N67A, K80A, K87P |  |  | N67A, K80A, K87P |  | 3 |
| R090 | I64C, K87C |  |  | I64C, K87C |  | 3 |
| R091 | K65C, K87C |  |  | K65C, K87C |  | 3 |
| R092 | L61C, Y86C |  |  | L61C, Y86C |  | 3 |
| R093 | S62C, Y86C |  |  | S62C, Y86C |  | 3 |
| R094 | L61C, V90C |  |  | L61C, V90C |  | 3 |
| R095 | N67C, L83C |  |  | N67C, L83C |  | 3 |
| R096 | N67C, K80C |  |  | N67C, K80C |  | 3 |
| R097 | S62C, K196C |  |  | S62C, K196C |  | 3 |
| R098 | N67C, V207C |  |  | N67C, V207C |  | 3 |
| R099 | L61C, L195C |  |  | L61C, L195C |  | 3 |
| R100 | I506D | I506D |  |  |  | 1 |
| R101 | I506E | I506E |  |  |  | 1 |
| R102 | F505H, I506D | F505H, I506D |  |  |  | 1 |
| R103 | F505H, I506E | F505H, I506E |  |  |  | 1 |
| R104 | F505K, I506D | F505K, I506D |  |  |  | 1 |
| R105 | F505K, I506E | F505K, I506E |  |  |  | 1 |
| R106 | I214P |  |  | I214P |  | 3 |
| R107 | S213P |  |  | S213P |  | 3 |
| R108 | Q210P |  |  | Q210P |  | 3 |
| R109 | S211P |  |  | S211P |  | 3 |
| R110 | N183P |  | N183P |  |  | 2 |
| R111 | S213P, I214P |  |  | S213P, I214P |  | 3 |
| R112 | Q210P, S211P |  |  | Q210P, S211P |  | 3 |
| R113 | S173P, T174P |  | S173P, T174P |  |  | 2 |
| R114 | L172P, S173P |  | L172P, S173P |  |  | 2 |
| R115 | S182P, N183P |  | S182P, N183P |  |  | 2 |
| R116 | L181P, S182P |  | L181P, S182P |  |  | 2 |
| R158 | K65C, K87C, F505H, I506D | F505H, I506D |  | K65C, K87C |  | 1+3 |
| R159 | K65C, K87C, F505K, I506E | F505K, I506E |  | K65C, K87C |  | 1+3 |
| R160 | K65C, K87C，S211P |  |  | K65C, K87C, S211P |  | 3 |
| R179 | K65C, K87C, A74F |  |  | K65C, K87C | A74F | 3+4 |
| R180 | K65C, K87C, A74W |  |  | K65C, K87C | A74W | 3+4 |
| R181 | K65C, K87C, A74Y |  |  | K65C, K87C | A74Y | 3+4 |
| R182 | K65C, K87C, N254Y |  |  | K65C, K87C | N254Y | 3+4 |
| R183 | K65C, K87C, A298K |  | A298K | K65C, K87C |  | 2+3 |
| R184 | K65C, K87C, A298Y |  | A298Y | K65C, K87C |  | 2+3 |
| R185 | K65C, K87C, G340F |  | G340F | K65C, K87C |  | 2+3 |
| R186 | K65C, K87C, G340Y |  | G340Y | K65C, K87C |  | 2+3 |
| R187 | K65C, K87C, D489F | D489F |  | K65C, K87C |  | 1+3 |
| R188 | K65C, K87C, S502F | S502F |  | K65C, K87C |  | 1+3 |
| R189 | K65C, K87C, S502W | S502W |  | K65C, K87C |  | 1+3 |
| R190 | K65C, K87C, S502Y | S502Y |  | K65C, K87C |  | 1+3 |
| R191 | S62L, K65C, K87C |  |  | S62L, K65C, K87C |  | 3 |
| R192 | S62P, K65C, K87C |  |  | S62P, K65C, K87C |  | 3 |
| R193 | K65C, K87C, S150L |  | S150L | K65C, K87C |  | 2+3 |
| R194 | K65C, K87C, G151L |  | G151L | K65C, K87C |  | 2+3 |
| R195 | K65C, K87C, S155I |  | S155I | K65C, K87C |  | 2+3 |
| R196 | K65C, K87C, S290P |  | S290P | K65C, K87C |  | 2+3 |
| R197 | K65C, K87C, A298I |  | A298I | K65C, K87C |  | 2+3 |
| R198 | K65C, K87C, A298L |  | A298L | K65C, K87C |  | 2+3 |
| R199 | K65C, K87C, T337W |  | T337W | K65C, K87C |  | 2+3 |
| R200 | K65C, K87C, G145I |  | G145I | K65C, K87C |  | 2+3 |
| R201 | K65C, K87C, L141C, N371C |  | L141C, N371C | K65C, K87C |  | 2+3 |
| R202 | K65C, K87C, V144C, M370C |  | V144C, M370C | K65C, K87C |  | 2+3 |
| R203 | K65C, K87C, L138C, Q354C |  | L138C, Q354C | K65C, K87C |  | 2+3 |
| R204 | K65C, K87C, L334C, I475C | L334C, I475C |  | K65C, K87C |  | 1+3 |
| R205 | K65C, K87C, F137C, P353C |  | F137C, P353C | K65C, K87C |  | 2+3 |
| R206 | K65C, K87C, I57C, S190C |  | I57C, S190C | K65C, K87C |  | 2+3 |
| R207 | K65C, K87C, S55C, L188C |  | S55C, L188C | K65C, K87C |  | 2+3 |
| R208 | K65C, K87C, H159C, I291C |  | H159C, I291C | K65C, K87C |  | 2+3 |
| R209 | K65C, K87C, Q34C, K470C | Q34C, K470C |  | K65C, K87C |  | 1+3 |
| R210 | K65C, K87C, Q34C, G471C | Q34C, G471C |  | K65C, K87C |  | 1+3 |
| R211 | K65C, K87C, S35C, P473C | S35C, P473C |  | K65C, K87C |  | 1+3 |
| R212 | K65C, K87C, I332C, F483C | I332C, F483C |  | K65C, K87C |  | 1+3 |
| R213 | K65C, K87C, I332C, P480C | I332C, P480C |  | K65C, K87C |  | 1+3 |
| R214 | K65C, K87C, M396C, F488C | M396C, F488C |  | K65C, K87C |  | 1+3 |
| R215 | K65C, K87C, K394C, S491C | K394C, S491C |  | K65C, K87C |  | 1+3 |
| R216 | K65C, K87C, T397C, E487C | T397C, E487C |  | K65C, K87C |  | 1+3 |
| R217 | K65C, K87C, T397C, D486C | T397C, D486C |  | K65C, K87C |  | 1+3 |
| R218 | K65C, K87C, T397C, S485C | T397C, S485C |  | K65C, K87C |  | 1+3 |
| R219 | K65C, K87C, F32C, Y468C | F32C, Y468C |  | K65C, K87C |  | 1+3 |
| R220 | K65C, K87C, E30C, S466C | E30C, S466C |  | K65C, K87C |  | 1+3 |
| R221 | K65C, K87C, Y33C, V469C | Y33C, V469C |  | K65C, K87C |  | 1+3 |
| R222 | K65C, K87C, E31C, L467C | E31C, L467C |  | K65C, K87C |  | 1+3 |
| R223 | K65C, K87C, T29C, K465C | T29C, K465C |  | K65C, K87C |  | 1+3 |
| R224 | K65C, K87C, Q26C, E463C | Q26C, E463C |  | K65C, K87C |  | 1+3 |
| R225 | K65C, K87C, K445C, E463C | K445C, E463C |  | K65C, K87C |  | 1+3 |
| R226 | K65C, K87C, S443C, S466C | S443C, S466C |  | K65C, K87C |  | 1+3 |
| R227 | K65C, K87C, T449C, Y458C | T449C, Y458C |  | K65C, K87C |  | 1+3 |
| R228 | K65C, K87C, K394W, D489F | K394W, D489F |  | K65C, K87C |  | 1+3 |
| R229 | K65C, K87C, K394F, D489F | K394F, D489F |  | K65C, K87C |  | 1+3 |
| R230 | K65C, K87C, K394Y, D489F | K394Y, D489F |  | K65C, K87C |  | 1+3 |
| R231 | K65C, K87C, K394L, D489F | K394L, D489F |  | K65C, K87C |  | 1+3 |
| R232 | K65C, K87C, K394W, D489L | K394W, D489L |  | K65C, K87C |  | 1+3 |
| R233 | K65C, K87C, K394F, D489L | K394F, D489L |  | K65C, K87C |  | 1+3 |
| R234 | K65C, K87C, K394Y, D489L | K394Y, D489L |  | K65C, K87C |  | 1+3 |
| R235 | K65C, K87C, K394L, D489L | K394L, D489L |  | K65C, K87C |  | 1+3 |
| R236 | K65C, K87C, L141R |  | L141R | K65C, K87C |  | 2+3 |
| R237 | K65C, K87C, L141K |  | L141K | K65C, K87C |  | 2+3 |
| R238 | K65C, K87C, Q354L, N371F |  | Q354L, N371F | K65C, K87C |  | 2+3 |
| R239 | K65C, K87C, Q354L, N371W |  | Q354L, N371W | K65C, K87C |  | 2+3 |
| R240 | K65C, K87C, Q354L, N371Y |  | Q354L, N371Y | K65C, K87C |  | 2+3 |
| R241 | K65C, K87C, S150L, Q302L |  | S150L, Q302L | K65C, K87C |  | 2+3 |
| R242 | K65C, K87C, S150L, Q302V |  | S150L, Q302V | K65C, K87C |  | 2+3 |
| R243 | K65C, K87C, S150L, G151L, Q302L |  | S150L, G151L, Q302L | K65C, K87C |  | 2+3 |
| R244 | K65C, K87C, S150L, G151L, Q302V |  | S150L, G151L, Q302V | K65C, K87C |  | 2+3 |
| R245 | K65C, K87C, I291E |  | I291E | K65C, K87C |  | 2+3 |
| R246 | K65C, K87C, I291D |  | I291D | K65C, K87C |  | 2+3 |
| R247 | K65C, K87C, S155V, S290V, A298L |  | S155V, S290V, A298L | K65C, K87C |  | 2+3 |
| R248 | K65C, K87C, S155V, S290A, A298L |  | S155V, S290A, A298L | K65C, K87C |  | 2+3 |
| R249 | K65C, K87C, Y458P | Y458P |  | K65C, K87C |  | 1+3 |
| R250 | K65C, K87C, V459P | V459P |  | K65C, K87C |  | 1+3 |
| R251 | K65C, K87C, N460P | N460P |  | K65C, K87C |  | 1+3 |
| R252 | K65C, K87C, K461P | K461P |  | K65C, K87C |  | 1+3 |
| R253 | K65C, K87C, Q462P | Q462P |  | K65C, K87C |  | 1+3 |
| R254 | K65C, K87C, E463P | E463P |  | K65C, K87C |  | 1+3 |
| R255 | K65C, K87C, V482P | V482P |  | K65C, K87C |  | 1+3 |
| R256 | K65C, K87C, F483P | F483P |  | K65C, K87C |  | 1+3 |
| R257 | K65C, K87C, K75C, I217C |  |  | K65C, K87C, K75C, I217C |  | 3 |
| R258 | K65C, K87C, A149C, Y458C |  | A149C, Y458C | K65C, K87C |  | 2+3 |
| R259 | K65C, K87C, G143C, V406C |  | G143C, V406C | K65C, K87C |  | 2+3 |
| R260 | K65C, K87C, G145C, I407C |  | G145C, I407C | K65C, K87C |  | 2+3 |
| R261 | K65C, K87C, F140C, S404C |  | F140C, S404C | K65C, K87C |  | 2+3 |
| R262 | K65C, K87C, A241C, Q279C |  |  | K65C, K87C | A241C, Q279C | 3+4 |
| R263 | K65C, K87C, A74K |  |  | K65C, K87C | A74K | 3+4 |
| R264 | K65C, K87C, A74R |  |  | K65C, K87C | A74R | 3+4 |
| R265 | K65C, K87C, A147V, T337F, R339W, Q354L, A355L, E356W, D368L, N371L, K394F, T400V, D489L, | K394F, T400V, D489L | A147V, T337F, R339W, 354L, A355L, 356W, D368L, N371L | K65C, K87C |  | 1+2+3 |
| R266 | K65C, K87C, A147V, Q354L, A355L, D368L, N371L |  | A147V, Q354L, A355L, D368L, N371L | K65C, K87C |  | 2+3 |
| R267 | K65C, K87C, T189V, A298F |  | T189V, A298F | K65C, K87C |  | 2+3 |
| R268 | K65C, K87C, L195K, N227E |  | N227E | K65C, K87C, L195K |  | 2+3 |
| R269 | K65C, K87C, S398F, D489L | S398F, D489L |  | K65C, K87C |  | 1+3 |
| R270 | K65C, K87C, N67L, I79L, V207I |  |  | K65C, K87C, N67L, I79L, V207I |  | 3 |
| R271 | K65C, K87C, S190F, R229Y, T253L, E256F |  | S190F, R229Y, E256F | K65C, K87C | T253L | 2+3+4 |
| R272 | K65C, K87C, S190L, R229W, T253L, E256F |  | S190L, R229W, E256F | K65C, K87C | T253L | 2+3+4 |
| R277 | K65C, K87C, F137C, T337C |  | F137C, T337C | K65C, K87C |  | 2+3 |
| R278 | K65C, K87C, F137C, D338C |  | F137C, D338C | K65C, K87C |  | 2+3 |
| R279 | K65C, K87C, F137C, R339C |  | F137C, R339C | K65C, K87C |  | 2+3 |
| R280 | K65C, K87C, L138C, D338C |  | L138C, D338C | K65C, K87C |  | 2+3 |
| R281 | K65C, K87C, L141C, L373C |  | L141C, L373C | K65C, K87C |  | 2+3 |
| R282 | K65C, K87C, L142C, P353C |  | L142C, P353C | K65C, K87C |  | 2+3 |
| R283 | K65C, K87C, L142C, N371C |  | L142C, N371C | K65C, K87C |  | 2+3 |
| R284 | K65C, K87C, G143C, N371C |  | G143C, N371C | K65C, K87C |  | 2+3 |
| R285 | K65C, K87C, V144C, N371C |  | V144C, N371C | K65C, K87C |  | 2+3 |
| R286 | K65C, K87C, G145C, M370C |  | G145C, M370C | K65C, K87C |  | 2+3 |
| R287 | K65C, K87C, S146C, M370C |  | S146C, M370C | K65C, K87C |  | 2+3 |
| R288 | K65C, K87C, S35C, I474C | S35C, I474C |  | K65C, K87C |  | 1+3 |
| R289 | K65C, K87C, K508C, S509C | K508C, S509C |  | K65C, K87C |  | 1+3 |
| R290 | K65C, K87C, L512C, L513C | L512C, L513C |  | K65C, K87C |  | 1+3 |
| R291 | K65C, K87C, V144C, M370C, I291E |  | V144C, M370C, I291E | K65C, K87C |  | 2+3 |
| R292 | K65C, K87C, V144C, M370C, I291D |  | V144C, M370C, I291D | K65C, K87C |  | 2+3 |
| R293 | V144C, M370C |  | V144C, M370C |  |  | 2 |
| R294 | K65C, K87C, V144C, M370C, A74F |  | V144C, M370C | K65C, K87C | A74F | 2+3+4 |
| R295 | K65C, K87C, V144C, M370C, A74W |  | V144C, M370C | K65C, K87C | A74W | 2+3+4 |
| R296 | K65C, K87C, V144C, M370C, A74Y |  | V144C, M370C | K65C, K87C | A74Y | 2+3+4 |
| R297 | K65C, K87C, V144C, M370C, F505K, I506E | F505K, I506E | V144C, M370C | K65C, K87C |  | 1+2+3 |
| R298 | K65C, K87C, V144C, M370C, S211P |  | V144C, M370C | K65C, K87C, S211P |  | 2+3 |
| R299 | K65C, K87C, V144C, M370C, A298Y |  | V144C, M370C, A298Y | K65C, K87C |  | 2+3 |
| R300 | K65C, K87C, V144C, M370C, T189V, A298F |  | V144C, M370C, T189V, A298F | K65C, K87C |  | 2+3 |
| R301 | K65C, K87C, V144C, M370C, N67L, I79L, V207I |  | V144C, M370C | K65C, K87C, N67L, I79L, V207I |  | 2+3 |
| R302 | N67A, K87V, V144C, M370C |  | V144C, M370C | N67A, K87V |  | 2+3 |
| R303 | K65C, K87C, V144C, M370C, K80W |  | V144C, M370C | K65C, K87C, K80W |  | 2+3 |
| R306 | K65C, K87C, V144C, M370C, K508C, S509C | K508C, S509C | V144C, M370C | K65C, K87C |  | 1+2+3 |
| R307 | K65C, K87C, V144C, M370C, L512C, L513C | L512C, L513C | V144C, M370C | K65C, K87C |  | 1+2+3 |
| R308 | K65C, K87C, V144C, M370C, A74Y, K508C, S509C | K508C, S509C | V144C, M370C | K65C, K87C | A74Y | 1+2+3+4 |
| R309 | K65C, K87C, V144C, M370C, A74Y, L512C, L513C | L512C, L513C | V144C, M370C | K65C, K87C | A74Y | 1+2+3+4 |
| R314 | K65C, K87C, V144C, M370C, A74Y, T522C, T523C | T522C, T523C | V144C, M370C | K65C, K87C | A74Y | 1+2+3+4 |
| R315 | K65C, K87C, V144C, M370C, A74Y, N515C, V516C | N515C, V516C | V144C, M370C | K65C, K87C | A74Y | 1+2+3+4 |
| R316 | K65C, K87C, V144C, M370C, A74Y, N515D | N515D | V144C, M370C | K65C, K87C | A74Y | 1+2+3+4 |
| R317 | K65C, K87C, V144C, M370C, A74Y, K520R | K520R | V144C, M370C | K65C, K87C | A74Y | 1+2+3+4 |
| R318 | K65C, K87C, V144C, M370C, A74Y, N515D, K520R | N515D, K520R | V144C, M370C | K65C, K87C | A74Y | 1+2+3+4 |
| R321 | K65C, K87C, V144C, M370C, A74Y, N183D |  | V144C, M370C, N183D | K65C, K87C | A74Y | 2+3+4 |
| R322 | K65C, K87C, V144C, M370C, A74Y, N183D, K427R |  | V144C, M370C, N183D, K427R | K65C, K87C | A74Y | 2+3+4 |
| R323 | K65C, K87C, V144C, M370C, A74Y, S404V |  | V144C, M370C, S404V | K65C, K87C | A74Y | 2+3+4 |
| R324 | K65C, K87C, V144C, M370C, A74Y, S404I |  | V144C, M370C, S404I | K65C, K87C | A74Y | 2+3+4 |
| R325 | K65C, K87C, V144C, M370C, A74Y, S404L |  | V144C, M370C, S404L | K65C, K87C | A74Y | 2+3+4 |
| R326 | K65C, K87C, V144C, M370C, A74Y, G143C, V406C |  | V144C, M370C, G143C, V406C | K65C, K87C | A74Y | 2+3+4 |
| R327 | K65C, K87C, V144C, M370C, A74Y, G145C, I407C |  | V144C, M370C, G145C, I407C | K65C, K87C | A74Y | 2+3+4 |
| R328 | K65C, K87C, V144C, M370C, A74Y, F140C, S404C |  | V144C, M370C, F140C, S404C | K65C, K87C | A74Y | 2+3+4 |
| R329 | K65C, K87C, V144C, M370C, A74Y, S150L |  | V144C, M370C, S150L | K65C, K87C | A74Y | 2+3+4 |
| R330 | K65C, K87C, V144C, M370C, A74Y, S150F |  | V144C, M370C, S150F | K65C, K87C | A74Y | 2+3+4 |
| R331 | K65C, K87C, V144C, M370C, A74Y, S150W |  | V144C, M370C, S150W | K65C, K87C | A74Y | 2+3+4 |
| R332 | K65C, K87C, V144C, M370C, A74Y, S150Y |  | V144C, M370C, S150Y | K65C, K87C | A74Y | 2+3+4 |
| R333 | K65C, K87C, V144C, M370C, A74Y |  | V144C, M370C | K65C, K87C | A74Y | 2+3+4 |
| R338 | N67A, A74Y, K87V, V144C, M370C |  | V144C, M370C | N67A, K87V | A74Y | 2+3+4 |
| R339 | K65C, K87C, A74Y, K508C, S509C | K508C, S509C |  | K65C, K87C | A74Y | 1+3+4 |
| R340 | K65C, K87C, V144C, M370C, N67A, A74Y |  | V144C, M370C | K65C, K87C, N67A | A74Y | 2+3+4 |
| R341 | N67V, A74Y, K87V, V144C, M370C |  | V144C, M370C | N67V, K87V | A74Y | 2+3+4 |
| R342 | N67V, A74Y, K80V, K87V, V144C, M370C |  | V144C, M370C | N67V, K80V, K87V | A74Y | 2+3+4 |
| R343 | N67V, A74Y, K80F, K87V, V144C, M370C |  | V144C, M370C | N67V, K80F, K87V | A74Y | 2+3+4 |
| R344 | N67V, A74Y, K80W, K87V, V144C, M370C |  | V144C, M370C | N67V, K80W, K87V | A74Y | 2+3+4 |
| R345 | N67V, A74Y, K80L, K87V, V144C, M370C |  | V144C, M370C | N67V, K80L, K87V | A74Y | 2+3+4 |
| R346 | N67V, A74Y, K80Y, K87L, V144C, M370C |  | V144C, M370C | N67V, K80Y, K87L | A74Y | 2+3+4 |
| R347 | K65C, K87C, V144C, M370C, N67V, A74Y |  | V144C, M370C | K65C, K87C, N67V | A74Y | 2+3+4 |
| R348 | K65C, K87C, V144C, M370C, N67V, A74Y, K80V |  | V144C, M370C | K65C, K87C, N67V, K80V | A74Y | 2+3+4 |
| R349 | K65C, K87C, V144C, M370C, N67V, A74Y, K80F |  | V144C, M370C | K65C, K87C, N67V, K80F | A74Y | 2+3+4 |
| R350 | K65C, K87C, V144C, M370C, N67V, A74Y, K80W |  | V144C, M370C | K65C, K87C, N67V, K80W | A74Y | 2+3+4 |
| R351 | K65C, K87C, V144C, M370C, N67V, A74Y, K80L |  | V144C, M370C | K65C, K87C, N67V, K80L | A74Y | 2+3+4 |
| R352 | K65C, K87C, V144C, M370C, N67V, A74Y, K80Y |  | V144C, M370C | K65C, K87C, N67V, K80Y | A74Y | 2+3+4 |

**Extended Data Table 2. Cryo-EM data processing and structure refinement statistics.**

|  | R296  (EMDB-81934)  (PDB-43KA) | Head38 in complex with CNR2292 and Motavizumab  (EMDB-81935)  (PDB-43KB) |
| --- | --- | --- |
| **Data collection** |  |  |
| Voltage (kV) | 300 | 300 |
| Microscope | FEI Titan Krios | FEI Titan Krios |
| Camera | Falcon4 | K3 (Gatan) |
| Magnification (calibrated) | 75,000X | 22,500x |
| Electron exposure (e^–^/Å^2^) | 50.43 | 60 |
| Exposure rate (e^–^/Å^2^/s) | 6.13 | 17.49 |
| Number of frames collected per micrograph | 32 | 32 |
| Automation software | EPU | SerialEM |
| Defocus range (μm) | –1.2 to –1.8 | –1.2 to –1.8 |
| Pixel size (Å) | 1.036 | 1.07 |
| **Overall map processing** |  |  |
| Micrographs used | 6,040 | 4,733 |
| Symmetry imposed | C3 | C1 |
| Initial particle images | 2,130,226 | 2,504,966 |
| Final particle images | 97,251 | 369,745 |
| Resolution at 0.143 FSC of masked reconstruction (Å) | 3.31 | 4.39 |
| Map sharpening B factor (Å^2^) | -134.4 | -258.4 |
| **Local map refinement** |  |  |
| Refinement package | Phenix v1.19 | Phenix v1.19 |
| Model composition |  |  |
| Non-hydrogen atoms | 10,455 | 5,150 |
| Protein residues | 1,356 | 664 |
| R.m.s. deviations |  |  |
| Bond lengths (Å) | 0.003 | 0.005 |
| Bond angles (°) | 0.551 | 1.075 |
| *B* factors (Å^2^) |  |  |
| Protein | 34.62 | 128.25 |
| Validation |  |  |
| MolProbity score | 1.75 | 1.93 |
| Clashscore | 6.49 | 8.95 |
| Poor rotamers (%) | 0 | 0 |
| Ramachandran plot |  |  |
| Favored (%) | 94.27 | 92.94 |
| Allowed (%) | 5.73 | 7.06 |
| Disallowed (%) | 0 | 0 |
| Cb outliers (%) | 0 | 0 |
| CaBLAM outliers (%) | 1.88 | 4.38 |

**Extended Data Table 3. Design and mutation information of epitope-enriched truncated constructs.**

| **Index** | **Signal peptide–antigen linker** | **N-terminal truncation segment** | **Inter-truncation linker** | **C-terminal truncation segment** | **Mutations within truncation segments** |
| --- | --- | --- | --- | --- | --- |
| 1 |  | 49-97 | GGSG | 145-308 | S155C, S290C, V207L, S190F |
| 2 |  | 49-97 | GGGSG | 145-308 | S155C, S290C, V207L, S190F |
| 3 |  | 49-97 | GGSGGS | 145-308 | S155C, S290C, V207L, S190F |
| 4 | GGS | 49-97 | GGSG | 145-308 | S155C, S290C, V207L, S190F |
| 5 | GGS | 49-97 | GGGSG | 145-308 | S155C, S290C, V207L, S190F |
| 6 | GGS | 49-97 | GGSGGS | 145-308 | S155C, S290C, V207L, S190F |
| 7 | GGS | 49-97 | GGSG | 145-308 | S155C, S290C, V207L, S190F, I59C, V192C |
| 8 | GGS | 49-97 | GGSG | 145-308 | S155C, S290C, V207L, S190F, E60C, D194C |
| 9 | GGS | 49-97 | GGSG | 145-308 | S155C, S290C, V207L, S190F, S55C, L188C |
| 10 | GGS | 49-97 | GGSG | 145-308 | S155C, S290C, V207L, S190F, I59C, L193C |
| 11 | GGS | 49-97 | GGSG | 145-308 | S155C, S290C, V207L, K176C, S190C |
| 12 | GGS | 49-97 | GGSG | 145-308 | S155C, S290C, V207L, S190F, A177C, T189C |
| 13 | GGS | 49-97 | GGSG | 145-308 | S155C, S290C, V207L, S190F, T58C, K191C |
| 14 | GGS | 49-97 | GGSG | 145-308 | S155C, S290C, V207L, S190F, L171C, K191C |
| 15 | GGS | 49-97 | GGSG | 145-308 | S155C, S290C, V207L, S190F, V56C, T189C |
| 16 | GGS | 49-97 | GGSG | 145-308 | S155C, S290C, V207L, I57C, S190C |
| 17 | GGS | 49-97 | GGSG | 145-308 | S155C, S290C, V207L, S190F, V56C, V187C |
| 18 | GGS | 49-97 | GGSG | 145-308 | S155C, S290C, V207L, S190F, T54C, G151C |
| 19 | GGS | 49-97 | GGGSG | 145-308 | S155C, S290C, V207L, S190F, I59C, V192C |
| 20 | GGS | 49-97 | GGGSG | 145-308 | S155C, S290C, V207L, S190F, E60C, D194C |
| 21 | GGS | 49-97 | GGGSG | 145-308 | S155C, S290C, V207L, S190F, S55C, L188C |
| 22 | GGS | 49-97 | GGGSG | 145-308 | S155C, S290C, V207L, S190F, I59C, L193C |
| 23 | GGS | 49-97 | GGGSG | 145-308 | S155C, S290C, V207L, K176C, S190C |
| 24 | GGS | 49-97 | GGGSG | 145-308 | S155C, S290C, V207L, S190F, A177C, T189C |
| 25 | GGS | 49-97 | GGGSG | 145-308 | S155C, S290C, V207L, S190F, T58C, K191C |
| 26 | GGS | 49-97 | GGGSG | 145-308 | S155C, S290C, V207L, S190F, L171C, K191C |
| 27 | GGS | 49-97 | GGGSG | 145-308 | S155C, S290C, V207L, S190F, V56C, T189C |
| 28 | GGS | 49-97 | GGGSG | 145-308 | S155C, S290C, V207L, I57C, S190C |
| 29 | GGS | 49-97 | GGGSG | 145-308 | S155C, S290C, V207L, S190F, V56C, V187C |
| 30 | GGS | 49-97 | GGGSG | 145-308 | S155C, S290C, V207L, S190F, T54C, G151C |
| 31 | GGS | 49-97 | GGSGGS | 145-308 | S155C, S290C, V207L, S190F, I59C, V192C |
| 32 | GGS | 49-97 | GGSGGS | 145-308 | S155C, S290C, V207L, S190F, E60C, D194C |
| 33 | GGS | 49-97 | GGSGGS | 145-308 | S155C, S290C, V207L, S190F, S55C, L188C |
| 34 | GGS | 49-97 | GGSGGS | 145-308 | S155C, S290C, V207L, S190F, I59C, L193C |
| 35 | GGS | 49-97 | GGSGGS | 145-308 | S155C, S290C, V207L, K176C, S190C |
| 36 | GGS | 49-97 | GGSGGS | 145-308 | S155C, S290C, V207L, S190F, A177C, T189C |
| 37 | GGS | 49-97 | GGSGGS | 145-308 | S155C, S290C, V207L, S190F, T58C, K191C |
| 38 | GGS | 49-97 | GGSGGS | 145-308 | S155C, S290C, V207L, S190F, L171C, K191C |
| 39 | GGS | 49-97 | GGSGGS | 145-308 | S155C, S290C, V207L, S190F, V56C, T189C |
| 40 | GGS | 49-97 | GGSGGS | 145-308 | S155C, S290C, V207L, I57C, S190C |
| 41 | GGS | 49-97 | GGSGGS | 145-308 | S155C, S290C, V207L, S190F, V56C, V187C |
| 42 | GGS | 49-97 | GGSGGS | 145-308 | S155C, S290C, V207L, S190F, T54C, G151C |
